# Mathematical models incorporating a multi-stage cell cycle replicate normally-hidden inherent synchronisation in cell proliferation

**DOI:** 10.1101/557702

**Authors:** Sean T. Vittadello, Scott W. McCue, Gency Gunasingh, Nikolas K. Haass, Matthew J. Simpson

## Abstract

We present a suite of experimental data showing that cell proliferation assays, prepared using standard methods thought to produce asynchronous cell populations, persistently exhibit inherent synchronisation. Our experiments use fluorescent cell cycle indicators to reveal the normally-hidden cell synchronisation by highlighting oscillatory subpopulations within the total cell population. These oscillatory subpopulations would never be observed without these cell cycle indicators. On the other hand, our experimental data show that the total cell population appears to grow exponentially, as in an asynchronous population. We reconcile these seemingly inconsistent observations by employing a multi-stage mathematical model of cell proliferation that can replicate the oscillatory subpopulations. Our study has important implications for understanding and improving experimental reproducibility. In particular, inherent synchronisation may affect the experimental reproducibility of studies aiming to investigate cell cycle-dependent mechanisms, including changes in migration and drug response.

## 1 Introduction

Cell proliferation is essential for a range of normal and pathological processes. Many different mathematical models of proliferation have been proposed [1–7]. It is often assumed that cells proliferate exponentially,

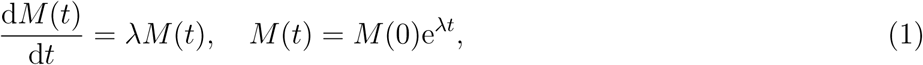

where *M* (*t*) is the number of cells at time *t* and *λ* > 0 is the proliferation rate.

The eukaryotic cell cycle consists of four phases in sequence, namely gap 1 (G1), synthesis (S), gap 2 (G2) and mitosis (M) (Figure 1(a)). A key assumption implicit in Equation (1) is that the cell population is asynchronous, meaning that the cells are distributed randomly among the cell cycle phases (Figure 1(b)), yielding a constant per capita growth rate, (1*/M* (*t*)) d*M* (*t*)*/*d*t* = *λ*. In contrast, a population of cells is synchronous if the cells are in the same cell cycle phase (Figure 1(c)), or partially synchronous if only a subpopulation of cells is synchronous (Figure 1(d)). In this case, the synchronous cells divide as a cohort in discrete stages producing a variable per capita growth rate. In addition to the implicit assumption of asynchronicity, classical exponential growth models, and generalisations thereof [8], do not account for subpopulations, and predict monotonic population growth.

**Figure 1:**
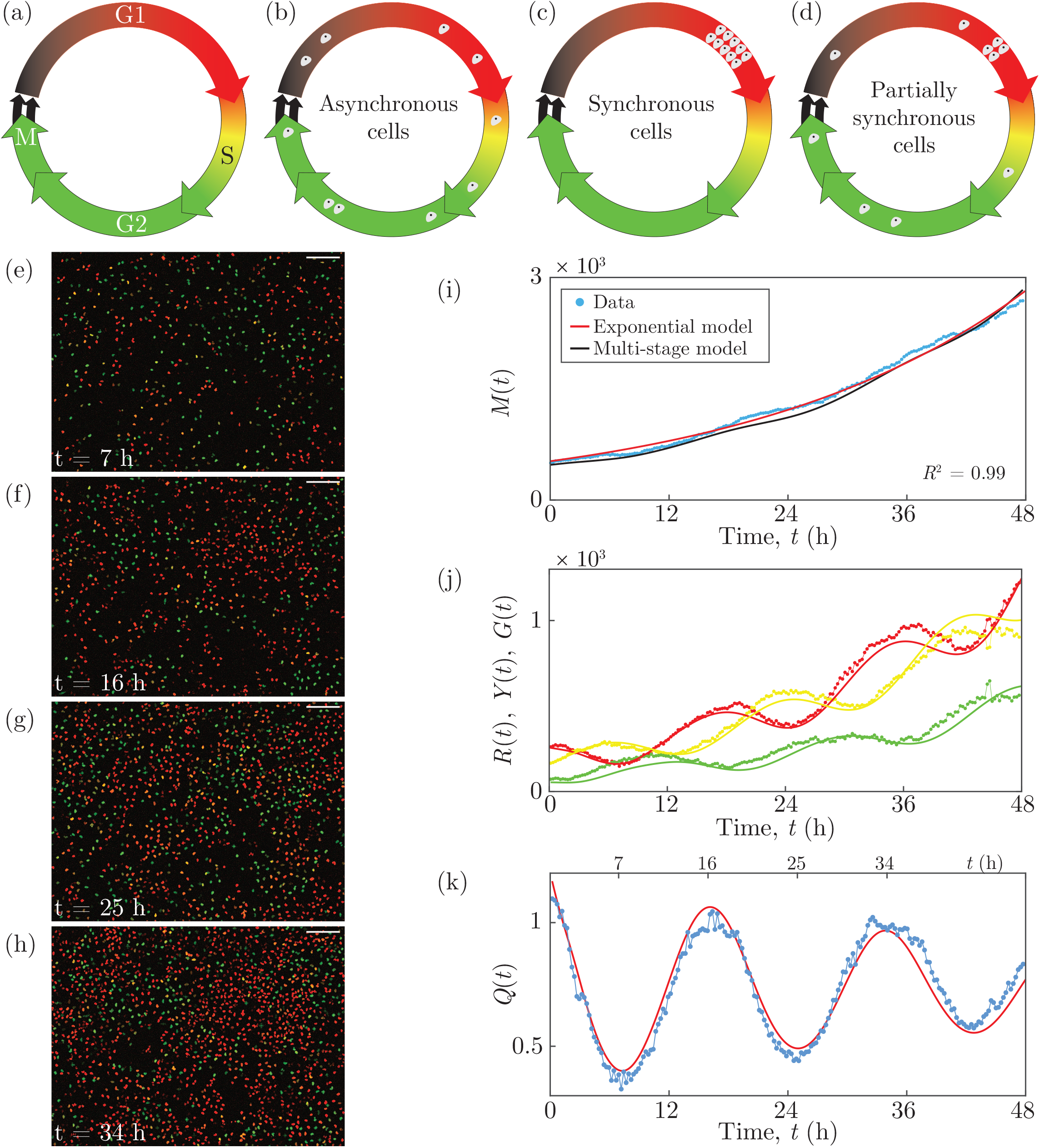
C8161 experimental data and multi-stage model solution. (a) The cell cycle, indicating the colour of FUCCI in each phase. (b)–(d) Asynchronous, synchronous and partially synchronous cells. (e)–(h) Images of a proliferation assay with FUCCI-C8161 cells. Scale bar 200 *µ*m. (i) *M* (*t*). Linear regression of ln *M* (*t*) versus *t* gives *R*^2^ = 0.99. (j) *R*(*t*), *Y* (*t*) and *G*(*t*). (k) *Q*(*t*). Experimental data are shown as discs and the model solutions as curves.

Here we provide new experimental data from two-dimensional cell proliferation assays in which the cell growth appears exponential as in Equation (1). Unexpectedly, however, we observe oscillatory subpopulations arising from a phenomenon we refer to as *inherent* synchronisation. We reveal the normally-hidden inherent synchronisation by identifying subpopulations based on cell cycle phase, employing fluorescent ubiquitination-based cell cycle indicator (FUCCI) [9]. FUCCI enables visualisation of the cell cycle of individual live cells via two sensors: when the cell is in G1 the nucleus fluoresces red, and when the cell is in S/G2/M the nucleus fluoresces green. During the G1/S transition, called early S (eS), both sensors fluoresce and the nucleus appears yellow (Figure 1(a)). We explain these seemingly inconsistent observations by applying a multi-stage mathematical model for cell proliferation.

Previous studies of cell synchronisation utilising FUCCI induce the synchronisation using methods including serum starvation, cell cycle-inhibiting drugs, environmental pH, or contact inhibition [10–15]. Our assays are prepared using a standard method [10] normally thought to produce asynchronous populations, and we take utmost care to ensure that there is no induced synchronisation in our cell cultures due to serum starvation, low pH, or contact inhibition (Supporting Information 1). Over three cell lines and four independent experiments, however, we consistently observe inherent synchronisation.

Neglecting synchronous subpopulations can have important implications for experiment reproducibility. For example, the accurate experimental evaluation of cell cycle-inhibiting drugs is highly dependent on the cell cycle distribution of the cell population [10,16]. In a partially synchronous population, the drug may have a delayed or advanced effect compared with an asynchronous population, depending on the cell cycle position of the synchronous cells. Generally, the presence of synchronisation may affect the reproducibility of experiments that investigate cell cycle-dependent mechanisms, such as changes in migration and drug response. Revealing any synchronisation with quantitative techniques like FUCCI will lead to a better understanding of these mechanisms.

## 2 Results

### 2.1 Experimental data

Our experimental data are time-series images from two-dimensional proliferation assays using three melanoma cell lines C8161, WM983C and 1205Lu [15, 17, 18], which have mean cell-cycle durations of approximately 18, 27 and 36 h, respectively [15]. Four independent experiments are performed for each cell line. Live-cell images are acquired at 15 minute intervals over 48 h.

Images from one position in a single well of a FUCCI-C8161 proliferation assay at 7, 16, 25 and 34 h show red, yellow or green nuclei corresponding to the phases G1, eS or S/G2/M (Figure 1(e)–(h)). We quantify the population growth by counting the total number of cells in each image (Supporting Information 1), to give *M* (*t*) at time *t* (Figure 1(i)). The total number of cells appears to grow exponentially over 48 h, supported by the best fit of Equation (1) (Supporting Information 1) since we have *R*^2^ = 0.99 from the linear regression of ln *M* (*t*) versus *t*. The temporal variations in the numbers of cells in the subpopulations *R*(*t*), *Y* (*t*) and *G*(*t*) with red, yellow or green nuclei (Figure 1(j)), where *M* (*t*) = *R*(*t*) + *Y* (*t*) + *G*(*t*), are oscillatory. In an asynchronous population, the subpopulations would exhibit monotone growth. The oscillations we observe, however, reveal that the cells are partially synchronous.

To explore the inherent synchronisation further, we group cells in eS and S/G2/M together, since eS is part of S, and consider the ratio *Q*(*t*) = *R*(*t*)*/* (*Y* (*t*) + *G*(*t*)) (Figure 1(k)). Synchronisation is clearly evident in the oscillatory nature of *Q*(*t*). Note the troughs at 7 and 25 h and the peaks at 16 and 34 h are separated by 18 h, which is the approximate cell cycle time for C8161. We can visualise the oscillations in these two subpopulations (Figure 1(e)–(h)), where the ratio of the number of red cells to the number of yellow and green cells is lower at 7 and 25 h and higher at 16 and 34 h. Equation (1) and related generalisations [8] cannot account for the oscillations in these subpopulations. Similar observations are made for further examples of this cell line, and the two additional cell lines (Supporting Information 1). We quantitatively confirm the presence of oscillations in *Q(t)*, arising from inherent synchronisation, for all 90 data sets by calculating the discrete Fourier transform of the *Q(t)* signal, and identifying the distinct dominant frequencies (Supporting Information 1). These results confirm that all 90 experimental replicates display oscillatory subpopulations that are inconsistent with traditional exponential and logistic growth models.

### 2.2 Multi-stage mathematical model

We employ a multi-stage model of cell proliferation [19] which can describe synchronous populations. The model assumes that the cell cycle durations follow a hypoexponential distribution, which consists of a series of independent exponential distributions with different rates. To apply this model we partition the cell cycle into *k* stages, *P*_*i*_ for *i* = 1, …, *k*, where the duration of each *P*_*i*_ is exponentially distributed with mean *µ*_*i*_. If *𝒯* is the mean cell cycle time then 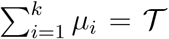. The stages *P*_*i*_ do not necessarily correspond to phases of the cell cycle, but instead are a mathematical device which allows control over the variance of cell-cycle phase durations in the multi-stage model, whereby more stages correspond to less variance in the phase durations for a cell population. If we let the transition rates be *λ*_*i*_ = 1*/µ*_*i*_ and consider the partitioned cell cycle 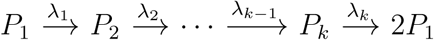, we arrive at a system of differential equations describing the mean population *M*_*i*_(*t*) in each stage [19]:

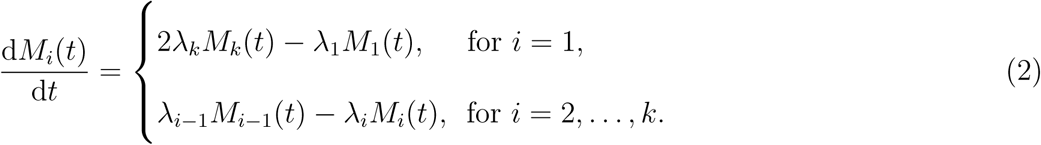

Note that 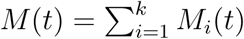. If *k* = 1, Equation (2) simplifies to Equation (1). Within the 48 h duration of our experiments, none of the cell lines exhibits contact inhibition of proliferation, consistent with the typical loss of contact inhibition in cancer cells [20]. Consequently, a carrying capacity is not incorporated into the model.

We solve Equation (2) numerically with the forward Euler method, and estimate the parameters by fitting the solution to our experimental data (Supporting Information 1). Using 18 stages for each of the three cell cycle phases described by FUCCI, giving *k* = 54, we obtain *M* (*t*) (Figure 1(i)), *R*(*t*), *Y* (*t*), *G*(*t*) (Figure 1(j)) and *Q*(*t*) (Figure 1(k)) which all correspond well with the experimental data. In particular, the multi-stage model replicates the oscillations in *R*(*t*), *Y* (*t*), *G*(*t*) and *Q*(*t*), a feature that is not possible with traditional exponential models. While the multi-stage model can replicate the oscillatory subpopulations, the model is unable to predict all features of the inherent synchronisation in a cell proliferation experiment due to variable initial conditions, as the inherent synchronisation is a stochastic phenomenon which likely arises from cell division and intercellular interactions. The model can, however, be used to predict general features of the inherent synchronisation of each cell line.

## 3 Conclusion

Our new experimental data demonstrate that cell populations may appear to grow exponentially despite subpopulations exhibiting oscillatory growth arising from normally-hidden inherent synchronisation. We use standard experimental methods thought to produce asynchronous populations; however, all of our proliferation assays exhibit inherent synchronisation. We use FUCCI to track cell-cycle progression, which is necessary to confirm cell synchronisation. As the standard exponential growth model cannot account for subpopulations with oscillating growth, we use a multi-stage mathematical model of cell proliferation to replicate oscillations in population growth. Our results are important because revealing any synchronisation will help to better understand cell cycle-dependent mechanisms, such as changes in migration and drug response. Without quantitative techniques like FUCCI to probe the cell cycle, synchronisation and its effects on experimental outcomes and reproducibility may remain hidden.

## 4 Author Contributions

All authors designed the research. STV performed the research. All authors contributed analytic tools and analysed the data. STV wrote the manuscript, and all authors approved the final version of the manuscript.

## 5 Acknowledgments

We thank the editor and four anonymous referees for helpful comments. NKH is a Cameron fellow of the Melanoma and Skin Cancer Research Institute, and is supported by the NHMRC (APP1084893). MJS is supported by the ARC (DP170100474).

