## Supplementary Material for "Mathematical models incorporating a multi-stage cell cycle replicate normally-hidden inherent synchronisation in cell proliferation"

*July 8, 2019*

**References** **44**

### 1 Experimental

Here we provide details of the experimental methodology that we use for our cell proliferation experiments involving three melanoma cell lines.

#### 1.1 Cell culture

The human melanoma cell lines C8161 (kindly provided by Mary Hendrix, Chicago, IL, USA), WM983C and 1205Lu (both kindly provided by Meenhard Herlyn, Philadelphia, PA, USA) were genotypically characterised [1–4], grown as described [5] (using 4% fetal bovine serum instead of 2%), and authenticated by STR fingerprinting (QIMR Berghofer Medical Research Institute, Herston, Australia).

We maintain the cell cultures to prevent any induced synchronisation from cell cycle arrest in G1 phase. In general, such induced synchronisation can occur through various experimental conditions, namely contact inhibition of proliferation at relatively high population densities [6], decreased pH of the growth medium due to the concentration of acidic cell-metabolites such as lactic acid [7], and reduced availability of nutrients such as serum [8]. We prevent induced synchronisation by passaging the cells every three days, and on the day prior to setting up an experiment, to maintain a subconfluent cell density and a fresh growth medium, so that the cell culture conditions are never such that they cause G1 arrest.

#### 1.2 Proliferation experiments

Cells are seeded from subconfluent culture flasks onto a 24-well plate at a density of $10^4$ cells cm$^{-2}$, with 2.5 ml of medium per well, which is 2.5 times the volume of the standard protocol. After incubating the plate for 24 h at 37°C with 5% $CO_2$, live-cell images are acquired at 15 minute intervals over 48 h at six different positions of the well. Four independent experiments are performed for each cell line.

Our preliminary experiments used a standard 1 ml of medium in each well, however the cells started to arrest in G1 around 48 hours after seeding, which is likely due to decreased pH of the medium from the lactic acid concentration. We therefore performed a large number of preliminary tests in an attempt to prevent the cells from arresting in G1 during the experiment, and we found that this is possible by increasing the volume of medium in each well to the reasonable maximum of 2.5 ml, given the volume of each well is 3 ml. The result of the increased volume of medium is that the cells do not begin to arrest in G1 until close to 72 hours following seeding, which provides us with almost 48 hours of imaging using cells that have minimal G1 arrest.

#### 1.3 Fluorescent ubiquitination-based cell cycle indicator (FUCCI)

To generate stable melanoma cell lines expressing the FUCCI constructs, mKO2-hCdt1 (30-120) and mAG-hGem (1-110) [9] were subcloned into a replication-defective, self-inactivating lentiviral expression vector system as previously described [5]. The lentivirus was produced by co-transfection of human embryonic kidney 293T cells. High-titer viral solutions for mKO2-hCdt1 (30/120) and mAG-hGem (1/110) were prepared and used for co-transduction into the melanoma cell lines, and subclones were generated by single cell sorting [10–12].

### 2 Image processing and analysis

The microscopy data consist of multi-channel time-series stacks which are processed and analysed automatically with Fiji/ImageJ and MATLAB as described below.

#### 2.1 Preprocessing

To maximise the accuracy in identifying particles, which in our case are cell nuclei, we enhance the quality of the microscopy images using ImageJ as follows.

1. Import the time-series stack with the Bio-Formats Importer plugin, splitting the red and green channels.
2. Apply five iterations of Subtract Background with rolling-ball radius of 5 pixels.
3. Apply Enhance Contrast with the Equalize Histogram option selected.
4. Apply the Gaussian Blur filter with sigma = 1.

#### 2.2 Segmentation

We now identify the particles in the processed images using ImageJ.

1. Apply Auto-thresholding using the Yen method, selecting the option to 'calculate the threshold for each image'.
2. The resulting binary images are then refined by applying:
   (a) Watershed;
   (b) Fill Holes;
   (c) Open, with iterations = 10 and count = 5;
   (d) Watershed.

#### 2.3 Analysis

For every image in the segmented binary time-series stacks we count the number of particles in each of the red and green channels using ImageJ. We then use MATLAB to determine which particles are yellow.

1. For each of the red and green channels, apply Analyze Particles in ImageJ with sizes in the range 5–$\infty$ pixels$^2$ and the option 'limit to threshold' selected. Output the stack position and the centroid of every particle in each channel.

2. We now need to determine which particles are red, yellow or green. A particle is red if it appears in the red channel, and there is no corresponding particle in the green channel. Similarly, a particle is green if it appears in the green channel, and there is no corresponding particle in the red channel. A particle is then yellow if it appears in both the red and green channels. Identifying whether a particle appears in both the red and green channels is complicated by the possible alteration of the shape of the particle during image processing. While we process every image in exactly the same way, the original microscopy images may have different signal-to-noise ratios between the red and green channels. Consequently, there may be a difference in the shape of a particle depending on the channel in which it is viewed, and thereby a difference in the centroid of the particle in each channel. We therefore use MATLAB to determine which particles are red, yellow or green, using the stack position and centroid of each particle, as follows.

   (a) We first find the yellow particles using the stack position and centroid of each particle, so choose a particle, in turn, from the red channel.

   (b) Search the green channel for a corresponding particle such that the Euclidean distance between the centroids of the two particles is not greater than 3 pixels, noting that the pixel size in our images is 1.8150 $\mu$m. This distance allows for a location error of the centroids of the red and green particles, whereby the centroids may be translated up to one pixel from the original centroid of the yellow particle in the unprocessed images. Placing the original yellow centroid at the centre of a $3 \times 3$

grid of pixels, the red and green centroids from the processed images may be located at any of the nine pixels in the grid.

(c) Once all of the yellow particles are found, the red particles are all of the particles in the red channel which are not yellow. Similarly, the green particles are all of the particles in the green channel which are not yellow.

### 3 Parameterisation of the exponential model

To estimate the parameters of the exponential model Equation (1) when fitting the model solution to the experimental data for the total number of cells, we use the the `fit` function and `exp1` model [13] in MATLAB. The parameter estimates, with 95% confidence intervals, are:

- **C8161 cell line - Figure 1(i)**

  $M(0) = 524.3$ $(515.1, 533.4)$ and $\lambda = 0.03504$ h$^{-1}$ $(0.03456, 0.03551)$.

- **C8161 cell line - Figure S2(e)**

  $M(0) = 386.4$ $(382.2, 390.5)$ and $\lambda = 0.0316$ h$^{-1}$ $(0.0313, 0.0319)$.

- **C8161 cell line - Figure S3(e)**

  $M(0) = 401$ $(393.8, 408.2)$ and $\lambda = 0.03573$ h$^{-1}$ $(0.03525, 0.03622)$.

- **WM983C cell line - Figure S4(e)**

  $M(0) = 247.7$ $(244.3, 251.2)$ and $\lambda = 0.02541$ h$^{-1}$ $(0.02501, 0.02581)$.

- **WM983C cell line - Figure S5(e)**

  $M(0) = 366.4$ $(362.8, 370)$ and $\lambda = 0.01917$ h$^{-1}$ $(0.01888, 0.01946)$.

- **WM983C cell line - Figure S6(e)**

  $M(0) = 158$ $(155.4, 160.7)$ and $\lambda = 0.0175$ h$^{-1}$ $(0.01699, 0.01801)$.

- **1205Lu cell line - Figure S7(e)**

  $M(0) = 215.9$ $(214.1, 217.7)$ and $\lambda = 0.01932$ h$^{-1}$ $(0.01907, 0.01958)$.

- **1205Lu cell line - Figure S8(e)**

  $M(0) = 249.1$ $(246.5, 251.7)$ and $\lambda = 0.01934$ h$^{-1}$ $(0.01903, 0.01965)$.

- **1205Lu cell line - Figure S9(e)**

  $M(0) = 266.6$ $(263.6, 269.6)$ and $\lambda = 0.01926$ h$^{-1}$ $(0.01893, 0.0196)$.

### 4 Parameterisation of the multi-stage mathematical model

Here we describe our methodology for estimating the parameters of the multi-stage mathematical model Equation (2) and fitting the model solution to the experimental data.

#### 4.1 Method for parameter estimation

The multi-stage model requires specification of the number of stages, the transition rates from each stage to the successive stage, and the initial population in each stage. In this work, we aim to achieve the best fit of the model to our data while keeping the number of model parameters with distinct values to a minimum.

We partition the phases G1, eS and S/G2/M into the same number of stages, $N$. The mean durations of the phases G1, eS and S/G2/M are denoted by $L_{\mathrm{r}}$, $L_{\mathrm{y}}$ and $L_{\mathrm{g}}$, respectively. The transition rates between successive stages are set equal within each phase to $N/L_{\mathrm{r}}$ in G1, $N/L_{\mathrm{y}}$ in eS, and $N/L_{\mathrm{g}}$ in S/G2/M. For each $i = 1, \dots, N$ we denote the mean number of cells at time $t$ in stage $i$ of G1 as $R_i(t)$, of eS as $Y_i(t)$, and of S/G2/M as $G_i(t)$. Therefore, $R(t) = \sum_{i=1}^{N} R_i(t)$, $Y(t) = \sum_{i=1}^{N} Y_i(t)$, and $G(t) = \sum_{i=1}^{N} G_i(t)$. The parameters that we need to estimate are the components of the vector

$$\mathbf{x} = [\, R_1(0) \dots R_N(0) \quad Y_1(0) \dots Y_N(0) \quad G_1(0) \dots G_N(0) \quad L_{\mathrm{r}} \quad L_{\mathrm{y}} \quad L_{\mathrm{g}} \,]. \tag{S1}$$

The parameters in Equation (S1) are either numbers of cells or phase durations, which are all non-negative, so we require our optimisation algorithm to accept bound constraints. To find estimates for these parameters we use the MATLAB nonlinear least-squares solver `lsqnonlin` [14] with the trust-region-reflective algorithm [15], which allows for bound constraints of the parameters. In the following, a dependent variable has the subscript 'model' or 'data' to distinguish between model and data values of the variable. With non-negative weights $w_2, \dots, w_7$, we define the vector objective function

$$\mathbf{F}(\mathbf{x}) = [\, \mathbf{f}_1(\mathbf{x}) \; w_2\mathbf{f}_2(\mathbf{x}) \; w_3\mathbf{f}_3(\mathbf{x}) \; w_4\mathbf{f}_4(\mathbf{x}) \; w_5\mathbf{f}_5(\mathbf{x}) \; w_6\mathbf{f}_6(\mathbf{x}) \; w_7\mathbf{f}_7(\mathbf{x}) \,] \tag{S2}$$

as the concatenation of the weight-scaled vectors

$$\mathbf{f}_1(\mathbf{x}) = \big[\, \big(Q_{\text{model}}(\mathbf{x}) - Q_{\text{data}}\big)(t_1) \quad \dots \quad \big(Q_{\text{model}}(\mathbf{x}) - Q_{\text{data}}\big)(t_n) \,\big], \tag{S3}$$

$$\mathbf{f}_2(\mathbf{x}) = \big[\, \big(R_{\text{model}}(\mathbf{x}) - R_{\text{data}}\big)(t_1) \quad \dots \quad \big(R_{\text{model}}(\mathbf{x}) - R_{\text{data}}\big)(t_n) \,\big], \tag{S4}$$

$$\mathbf{f}_3(\mathbf{x}) = \big[\, \big(Y_{\text{model}}(\mathbf{x}) - Y_{\text{data}}\big)(t_1) \quad \dots \quad \big(Y_{\text{model}}(\mathbf{x}) - Y_{\text{data}}\big)(t_n) \,\big], \tag{S5}$$

$$\mathbf{f}_4(\mathbf{x}) = \big[\, \big(G_{\text{model}}(\mathbf{x}) - G_{\text{data}}\big)(t_1) \quad \dots \quad \big(G_{\text{model}}(\mathbf{x}) - G_{\text{data}}\big)(t_n) \,\big], \tag{S6}$$

$$\mathbf{f}_5(\mathbf{x}) = \big[\, \big(G_{\text{model}}(\mathbf{x}) - G_{\text{data}}\big)(t_n) \,\big], \tag{S7}$$

$$\mathbf{f}_6(\mathbf{x}) = \big[\, \mathcal{T} - L_{\text{r}} - L_{\text{y}} - L_{\text{g}} \,\big], \tag{S8}$$

$$\mathbf{f}_7(\mathbf{x}) = \big[\, \big(M_{\text{model}}(\mathbf{x}) - M_{\text{data}}\big)(t_1) \quad \dots \quad \big(M_{\text{model}}(\mathbf{x}) - M_{\text{data}}\big)(t_n) \,\big], \tag{S9}$$

where:

1. $Q_{\text{model}}(\mathbf{x})$ is the ratio of the number of cells in G1 to the number of cells in eS/S/G2/M, from the model solution;
2. $Q_{\text{data}}$ is the ratio of the number of cells in G1 to the number of cells in eS/S/G2/M, from the data;
3. $R_{\text{model}}(\mathbf{x})$, $Y_{\text{model}}(\mathbf{x})$ and $G_{\text{model}}(\mathbf{x})$ are the subpopulations of cells in G1, eS and S/G2/M, respectively, from the model solution;
4. $R_{\text{data}}$, $Y_{\text{data}}$ and $G_{\text{data}}$ are the subpopulations of cells in G1, eS and S/G2/M, respectively, from the data;
5. $M_{\text{model}}(\mathbf{x})$ is the total cell population, from the model solution;
6. $M_{\text{data}}$ is the total cell population, from the data;
7. $t_1 < \cdots < t_n$ are the data time points over 48 hours;
8. $\mathcal{T}$ is the cell cycle time.

The vector $\mathbf{f}_1$ is used to fit the model to the ratio data, and the vectors $\mathbf{f}_2, \ldots, \mathbf{f}_4$ are used to fit the model to the three subpopulations corresponding to G1, eS and S/G2/M. The vector $\mathbf{f}_5$ fits the model to the S/G2/M subpopulation data at the final time point, and is only required if the cells are starting to arrest in G1 near the end of the experiment due to the decreased pH of the growth medium. The vector $\mathbf{f}_6$ constrains the estimated phase durations to sum to the expected cell cycle time, and is generally required only when there are an insufficient number of oscillations in $Q_{\text{data}}(t)$ to bound the estimated cell-phase durations to physically realistic values. The vector $\mathbf{f}_7$ is used to fit the model to the total population data, and is often not required as a good fit usually follows from fitting to the subpopulations.

Note that the weights in the objective function Equation (S2) are specified prior to optimising the estimates of the parameters in Equation (S1). The weights differ between data sets in order to obtain the best fit of the multi-stage model to each data set.

#### 4.2 Specific estimated parameters

Here we provide a summary of the estimated parameters of the multi-stage model Equation (2) corresponding to Figure 1, along with additional data sets from the C8161 (Figures S2–S3), WM983C (Figures S4–S6) and 1205Lu (Figure S7–S9) cell lines. The correspondence between these figures and the data sets in Figures S10–S15 is:

- Figure 1 - First plot in Well 1 of Experiment 2, Figures S10 and S11;
- Figure S2 - Second plot in Well 1 of Experiment 1, Figures S10 and S11;
- Figure S3 - First plot in Well 1 of Experiment 3, Figures S10 and S11;
- Figure S4 - Second plot in Well 1 of Experiment 4, Figures S12 and S13;
- Figure S5 - Fourth plot in Well 1 of Experiment 1, Figures S12 and S13;
- Figure S6 - Third plot in Well 1 of Experiment 2, Figures S12 and S13;

- Figure S7 - First plot in Well 1 of Experiment 4, Figures S14 and S15;
- Figure S8 - Fourth plot in Well 1 Experiment 4, Figures S14 and S15;
- Figure S9 - Fifth plot in Well 2 of Experiment 4, Figures S14 and S15.

###### 4.2.1 C8161 cell line - Figure 1

The experimentally-determined mean cell cycle time for C8161 is approximately $\mathcal{T} = 18$ h [10]. We partition each cell cycle phase into $N = 18$ stages, giving a total of $k = 54$ stages for the complete cell cycle. In each phase we set the first half of the stages, totalling 9 stages, to have equal numbers of cells, and the second half of the stages to have equal numbers of cells. We therefore only require a total of six distinct population parameters.

The vector objective function is $\mathbf{F}(\mathbf{x}) = [\,\mathbf{f}_1(\mathbf{x})\ \ 10^{-4}\mathbf{f}_2(\mathbf{x})\ \ 10^{-4}\mathbf{f}_3(\mathbf{x})\ \ 10^{-4}\mathbf{f}_4(\mathbf{x})\ \ 10^{-3}\mathbf{f}_5(\mathbf{x})\,]$. Starting with the parameters $R_i(0) = Y_i(0) = G_i(0) = 0.5$ for $i = 1, \ldots, N$, and $L_\mathrm{r} = L_\mathrm{y} = L_\mathrm{g} = 6$, we obtain the parameterisation

$$R_i(0) = \begin{cases} 15.48 & \text{for } i = 1, \ldots, 9, \\ 12.93 & \text{for } i = 10, \ldots, 18, \end{cases} \qquad Y_i(0) = \begin{cases} 12.94 & \text{for } i = 1, \ldots, 9, \\ 5.41 & \text{for } i = 10, \ldots, 18, \end{cases}$$
$$G_i(0) = \begin{cases} 2.51 & \text{for } i = 1, \ldots, 9, \\ 3.48 & \text{for } i = 10, \ldots, 18, \end{cases} \qquad \begin{aligned} L_\mathrm{r} &= 6.14 \text{ h}, \\ L_\mathrm{y} &= 7.43 \text{ h}, \\ L_\mathrm{g} &= 4.42 \text{ h}. \end{aligned} \tag{S10}$$

All parameter estimates given in this document are presented to two decimal places. Note that $L_\mathrm{r} + L_\mathrm{y} + L_\mathrm{g} = 17.99$ h, in good agreement with the observed cell cycle time of 18 h.

##### 4.2.2 C8161 cell line, different numbers of stages - Figure S1

In Figure S1 we compare solutions of the multi-stage model for $N = 2$, 6, 10 and 14 stages per phase, with the ratio $Q_{\text{data}}$. In fitting the model solution we use the same parameters as for Figure 1, except the number

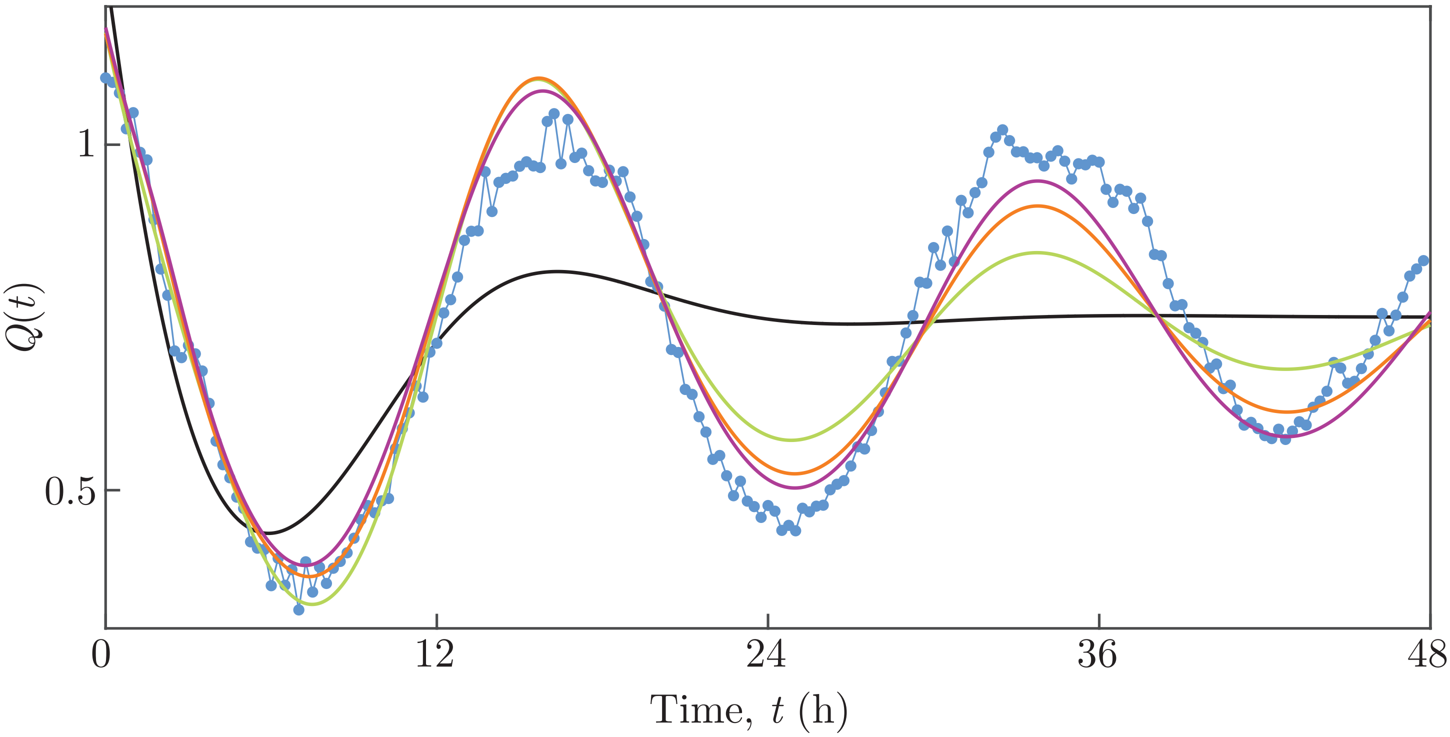

**Figure S1: $Q(t)$ for C8161 experimental data and multi-stage model solutions with different numbers of stages.** Experimental data are shown as discs and the model solutions as curves. The model solutions with 2, 6, 10 and 14 stages are the black, green, orange and purple curves, respectively.

of stages differ. For each number of stages, in each phase we set the first half of the stages to have equal numbers of cells and the second half of the stages to have equal numbers of cells, so that we therefore only require a total of 6 distinct population parameters. Starting with the parameters $R_i(0) = Y_i(0) = G_i(0) = 0.5$

for $i = 1, \ldots, N$, and $L_\mathrm{r} = L_\mathrm{y} = L_\mathrm{g} = 6$, the parameterisation for $N = 14$ is

$$R_i(0) = \begin{cases} 17.89 & \text{for } i = 1, \ldots, 9, \\ 18.62 & \text{for } i = 10, \ldots, 18, \end{cases} \qquad Y_i(0) = \begin{cases} 16.28 & \text{for } i = 1, \ldots, 9, \\ 6.77 & \text{for } i = 10, \ldots, 18, \end{cases}$$
$$L_\mathrm{r} = 6.17 \text{ h}, \tag{S11}$$
$$G_i(0) = \begin{cases} 2.39 & \text{for } i = 1, \ldots, 9, \\ 5.78 & \text{for } i = 10, \ldots, 18, \end{cases} \qquad \begin{aligned} L_\mathrm{y} &= 7.27 \text{ h}, \\ L_\mathrm{g} &= 4.64 \text{ h}, \end{aligned}$$

the parameterisation for $N = 10$ is

$$R_i(0) = \begin{cases} 16.60 & \text{for } i = 1, \ldots, 9, \\ 33.58 & \text{for } i = 10, \ldots, 18, \end{cases} \qquad Y_i(0) = \begin{cases} 20.81 & \text{for } i = 1, \ldots, 9, \\ 9.61 & \text{for } i = 10, \ldots, 18, \end{cases}$$
$$L_\mathrm{r} = 6.22 \text{ h}, \tag{S12}$$
$$G_i(0) = \begin{cases} 0 & \text{for } i = 1, \ldots, 9, \\ 12.81 & \text{for } i = 10, \ldots, 18, \end{cases} \qquad \begin{aligned} L_\mathrm{y} &= 7.06 \text{ h}, \\ L_\mathrm{g} &= 4.92 \text{ h}, \end{aligned}$$

the parameterisation for $N = 6$ is

$$R_i(0) = \begin{cases} 16.42 & \text{for } i = 1, \ldots, 9, \\ 66.47 & \text{for } i = 10, \ldots, 18, \end{cases} \qquad Y_i(0) = \begin{cases} 44.97 & \text{for } i = 1, \ldots, 9, \\ 0 & \text{for } i = 10, \ldots, 18, \end{cases}$$
$$L_\mathrm{r} = 6.28 \text{ h}, \tag{S13}$$
$$G_i(0) = \begin{cases} 0 & \text{for } i = 1, \ldots, 9, \\ 26.34 & \text{for } i = 10, \ldots, 18, \end{cases} \qquad \begin{aligned} L_\mathrm{y} &= 6.81 \text{ h}, \\ L_\mathrm{g} &= 5.21 \text{ h}, \end{aligned}$$

and the parameterisation for $N = 2$ is

$$R_i(0) = \begin{cases} 195.22 & \text{for } i = 1, \ldots, 9, \\ 107.41 & \text{for } i = 10, \ldots, 18, \end{cases} \qquad Y_i(0) = \begin{cases} 241.05 & \text{for } i = 1, \ldots, 9, \\ 0 & \text{for } i = 10, \ldots, 18, \end{cases}$$

$$L_{\mathrm{r}} = 6.91 \text{ h}, \tag{S14}$$

$$G_i(0) = \begin{cases} 0 & \text{for } i = 1, \ldots, 9, \\ 0 & \text{for } i = 10, \ldots, 18, \end{cases} \qquad L_{\mathrm{y}} = 7.11 \text{ h}, \quad L_{\mathrm{g}} = 5.79 \text{ h}.$$

The corresponding solutions of the multi-stage model are shown in Figure S1.

Considering parameterisations of the model whereby in each phase we set the first half of the stages to have equal numbers of cells and the second half of the stages to have equal numbers of cells, the oscillations decay at a faster rate for a smaller number of stages per phase. A higher number of stages produces a hypoexponential distribution with lower variance, resulting in oscillations which are sustained for longer. Consequently, fewer than 18 stages per phase results in a model solution with a poorer fit.

###### 4.2.3 C8161 cell line - Figure S2

The experimentally-determined mean cell cycle time for C8161 is approximately $\mathcal{T} = 18$ h [10]. We partition each cell cycle phase into $N = 40$ stages, giving a total of $k = 120$ stages for the complete cell cycle. In each phase we set the first half of the stages, totalling 20 stages, to have equal numbers of cells, and the second half of the stages to have equal numbers of cells. We therefore only require a total of six distinct population parameters.

The vector objective function is $\mathbf{F}(\mathbf{x}) = [\, \mathbf{f}_1(\mathbf{x}) \;\; 10^{-3}\mathbf{f}_2(\mathbf{x}) \;\; 10^{-3}\mathbf{f}_3(\mathbf{x}) \;\; 10^{-3}\mathbf{f}_4(\mathbf{x}) \,]$. Trialling different parameters chosen randomly and uniformly from $(0.1, 1)$ for $R_i(0)$, $Y_i(0)$ and $G_i(0)$, and from $(4, 8)$ for $L_{\mathrm{r}}$, $L_{\mathrm{y}}$ and

$L_\mathrm{g}$, we obtain the parameterisation

$$R_i(0) = \begin{cases} 2.56 & \text{for } i = 1, \ldots, 20, \\ 1.99 & \text{for } i = 21, \ldots, 40, \end{cases} \qquad Y_i(0) = \begin{cases} 5.37 & \text{for } i = 1, \ldots, 20, \\ 4.13 & \text{for } i = 21, \ldots, 40, \end{cases}$$
$$L_\mathrm{r} = 5.47 \text{ h}, \tag{S15}$$
$$G_i(0) = \begin{cases} 0.90 & \text{for } i = 1, \ldots, 20, \\ 0.98 & \text{for } i = 21, \ldots, 40, \end{cases} \qquad L_\mathrm{y} = 8.67 \text{ h}, \quad L_\mathrm{g} = 4.57 \text{ h}.$$

Note that $L_\mathrm{r} + L_\mathrm{y} + L_\mathrm{g} = 18.71$ h, in good agreement with the observed cell cycle time of 18 h.

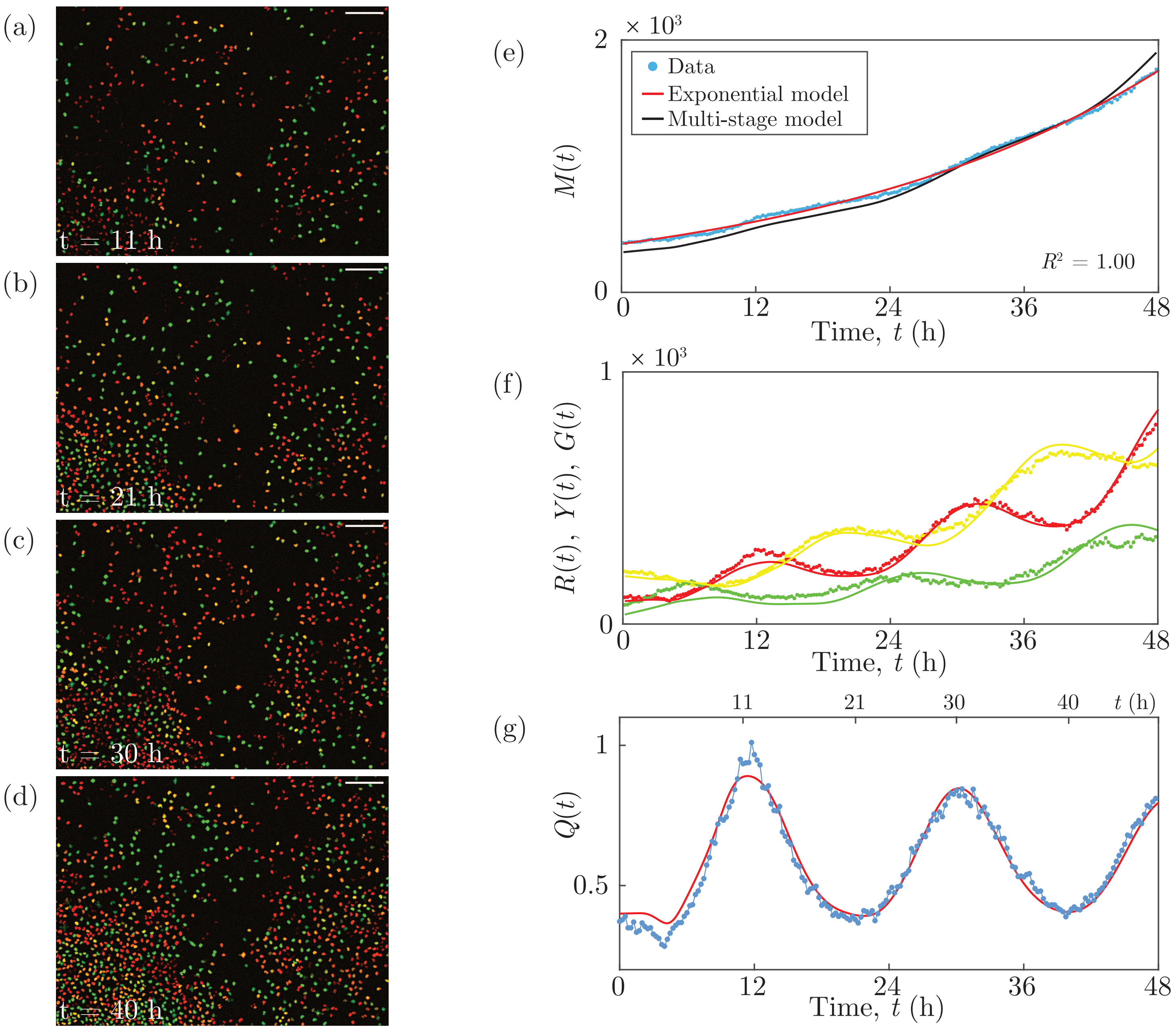

**Figure S2: C8161 experimental data and multi-stage model solution.** (a)–(d) Images of a proliferation assay with FUCCI-C8161 cells. Scale bar 200 $\mu$m. (e) $M(t)$. Linear regression of $\ln M(t)$ versus $t$ provides $R^2$. (f) $R(t)$, $Y(t)$ and $G(t)$. (g) $Q(t)$. Experimental data are shown as discs and the model solutions as curves.

###### 4.2.4 C8161 cell line - Figure S3

The experimentally-determined mean cell cycle time for C8161 is approximately $\mathcal{T} = 18$ h [10]. We partition each cell cycle phase into $N = 40$ stages, giving a total of $k = 120$ stages for the complete cell cycle. In each phase we set the first half of the stages, totalling 20 stages, to have equal numbers of cells, and the second half of the stages to have equal numbers of cells. We therefore only require a total of six distinct population parameters.

The vector objective function is $\mathbf{F}(\mathbf{x}) = [\,\mathbf{f}_1(\mathbf{x})\ \ 10^{-4}\mathbf{f}_2(\mathbf{x})\ \ 10^{-4}\mathbf{f}_3(\mathbf{x})\ \ 10^{-4}\mathbf{f}_4(\mathbf{x})\ \ 10^{-3}\mathbf{f}_5(\mathbf{x})\,]$. Trialling different parameters chosen randomly and uniformly from $(0.1, 1)$ for $R_i(0)$, $Y_i(0)$ and $G_i(0)$, and from $(4, 8)$ for $L_\mathrm{r}$, $L_\mathrm{y}$ and $L_\mathrm{g}$, we obtain the parameterisation

$$
\begin{aligned}
R_i(0) &= \begin{cases} 3.21 & \text{for } i = 1, \ldots, 20, \\ 4.44 & \text{for } i = 21, \ldots, 40, \end{cases} \qquad
Y_i(0) = \begin{cases} 5.03 & \text{for } i = 1, \ldots, 20, \\ 2.59 & \text{for } i = 21, \ldots, 40, \end{cases} \\
& \qquad\qquad\qquad\qquad\qquad\qquad\quad L_\mathrm{r} = 6.29 \text{ h}, \\
G_i(0) &= \begin{cases} 1.31 & \text{for } i = 1, \ldots, 20, \\ 1.34 & \text{for } i = 21, \ldots, 40, \end{cases} \qquad
\begin{aligned} L_\mathrm{y} &= 7.30 \text{ h}, \\ L_\mathrm{g} &= 4.47 \text{ h}. \end{aligned}
\end{aligned}
\tag{S16}
$$

Note that $L_\mathrm{r} + L_\mathrm{y} + L_\mathrm{g} = 18.06$ h, in good agreement with the observed cell cycle time of 18 h.

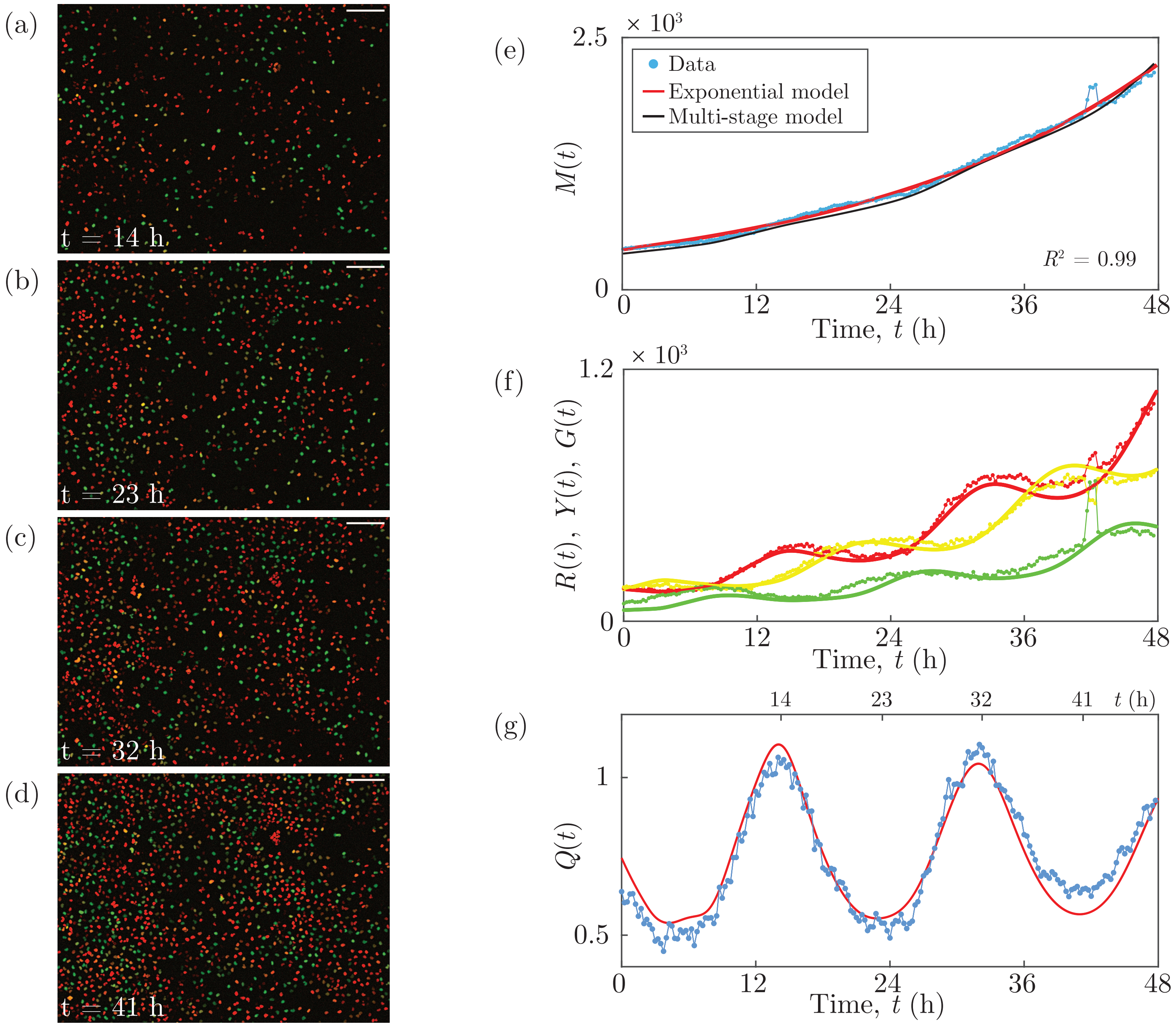

**Figure S3: C8161 experimental data and multi-stage model solution.** (a)–(d) Images of a proliferation assay with FUCCI-C8161 cells. Scale bar 200 $\mu$m. (e) $M(t)$. Linear regression of $\ln M(t)$ versus $t$ provides $R^2$. (f) $R(t)$, $Y(t)$ and $G(t)$. (g) $Q(t)$. Experimental data are shown as discs and the model solutions as curves.

###### 4.2.5 WM983C cell line - Figure S4

The experimentally-determined mean cell cycle time for WM983C is $\mathcal{T} = 27$ h [10]. We partition each cell cycle phase into $N = 10$ stages, giving a total of $k = 30$ stages for the complete cell cycle. From the start of each phase we set every 2 successive stages to have equal numbers of cells. We therefore only require a total of 15 distinct population parameters.

The vector objective function is $\mathbf{F}(\mathbf{x}) = [\,\mathbf{f}_1(\mathbf{x})\ \ 10^{-3}\mathbf{f}_2(\mathbf{x})\ \ 10^{-3}\mathbf{f}_3(\mathbf{x})\ \ 10^{-3}\mathbf{f}_4(\mathbf{x})\ \ 10\,\mathbf{f}_6(\mathbf{x})\,]$. Starting with the parameters

$$
\begin{aligned}
R_i(0) &= \begin{cases} 0 \quad \text{for } i = 1, 2, \\ 0 \quad \text{for } i = 3, 4, \\ 29.23 \quad \text{for } i = 5, 6, \\ 19.66 \quad \text{for } i = 7, 8, \\ 0 \quad \text{for } i = 9, 10, \end{cases} \qquad
Y_i(0) = \begin{cases} 33.02 \quad \text{for } i = 1, 2, \\ 0 \quad \text{for } i = 3, 4, \\ 0 \quad \text{for } i = 5, 6, \\ 0 \quad \text{for } i = 7, 8, \\ 0 \quad \text{for } i = 9, 10, \end{cases} \\
G_i(0) &= \begin{cases} 11.04 \quad \text{for } i = 1, 2, \\ 0 \quad \text{for } i = 3, 4, \\ 4.29 \quad \text{for } i = 5, 6, \\ 24.41 \quad \text{for } i = 7, 8, \\ 0 \quad \text{for } i = 9, 10, \end{cases} \qquad
\begin{aligned} L_{\mathrm{r}} &= 10.28 \text{ h}, \\ L_{\mathrm{y}} &= 3.87 \text{ h}, \\ L_{\mathrm{g}} &= 12.85 \text{ h}, \end{aligned}
\end{aligned}
\tag{S17}
$$

we obtain the same parameterisation Equation (S17). Note that $L_{\mathrm{r}} + L_{\mathrm{y}} + L_{\mathrm{g}} = 27.00$ h, in good agreement with the observed cell cycle time of 27 h.

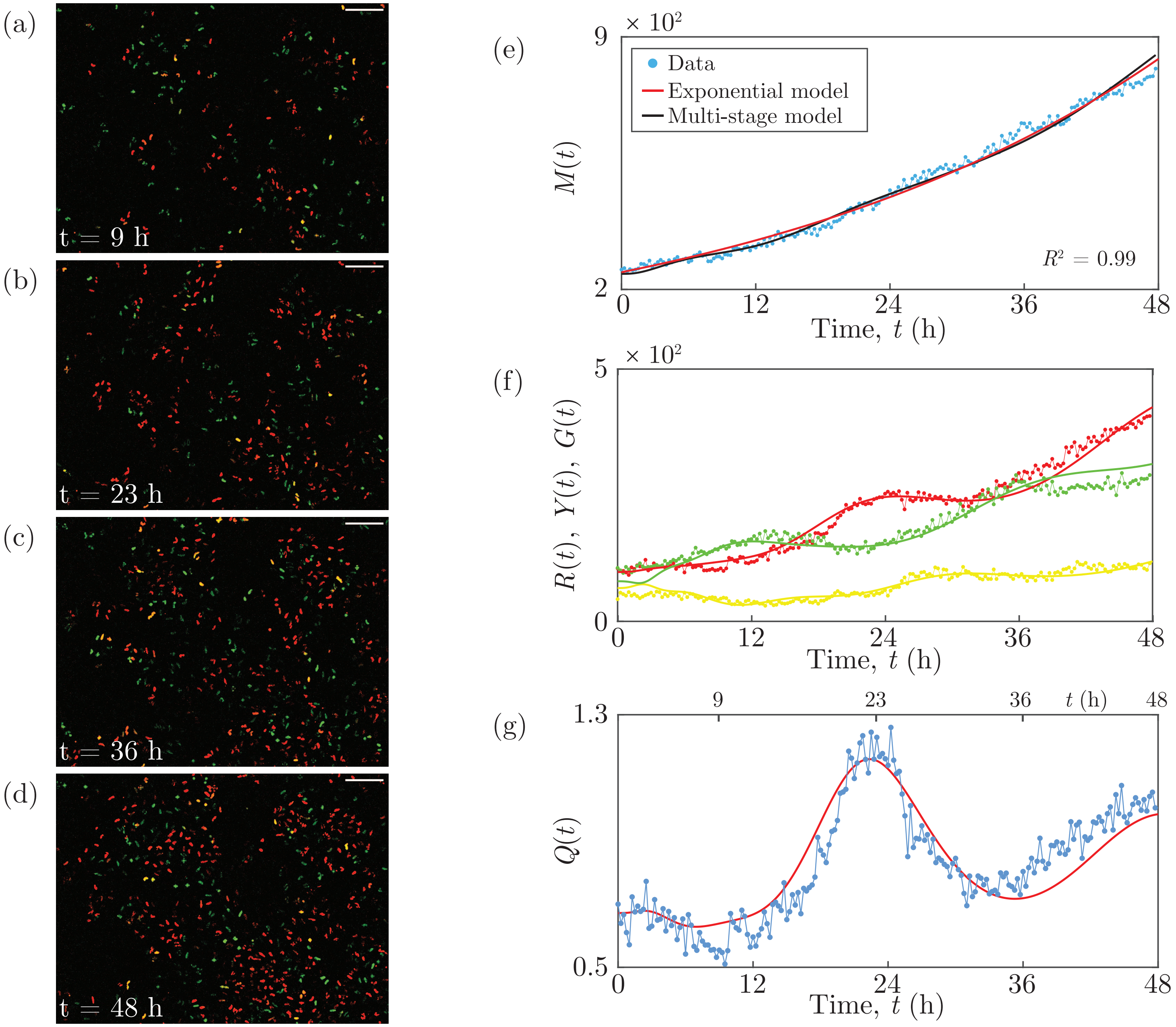

**Figure S4: WM983C experimental data and multi-stage model solution.** (a)–(d) Images of a proliferation assay with FUCCI-WM983C cells. Scale bar 200 $\mu$m. (e) $M(t)$. Linear regression of $\ln M(t)$ versus $t$ provides $R^2$. (f) $R(t)$, $Y(t)$ and $G(t)$. (g) $Q(t)$. Experimental data are shown as discs and the model solutions as curves.

###### 4.2.6 WM983C cell line - Figure S5

The experimentally-determined mean cell cycle time for WM983C is $\mathcal{T} = 27$ h [10]. We partition each cell cycle phase into $N = 10$ stages, giving a total of $k = 30$ stages for the complete cell cycle. From the start of each phase we set every 5 successive stages to have equal numbers of cells. We therefore only require a total of 6 distinct population parameters.

The vector objective function is $\mathbf{F}(\mathbf{x}) = [\,\mathbf{f}_1(\mathbf{x})\ \ 10^{-3}\mathbf{f}_2(\mathbf{x})\ \ 10^{-3}\mathbf{f}_3(\mathbf{x})\ \ 10^{-3}\mathbf{f}_4(\mathbf{x})\ \ 10^{-3}\mathbf{f}_7(\mathbf{x})\,]$. Trialling different parameters chosen randomly and uniformly from $(0.1, 1)$ for $R_i(0)$, $Y_i(0)$ and $G_i(0)$, and from $(4, 20)$ for $L_\mathrm{r}$, $L_\mathrm{y}$ and $L_\mathrm{g}$, we obtain the parameterisation

$$
\begin{aligned}
R_i(0) &= \begin{cases} 12.82 & \text{for } i = 1, \ldots, 5, \\ 13.13 & \text{for } i = 6, \ldots, 10, \end{cases} \qquad
Y_i(0) = \begin{cases} 7.86 & \text{for } i = 1, \ldots, 5, \\ 8.47 & \text{for } i = 6, \ldots, 10, \end{cases} \\
& \qquad\qquad\qquad\qquad\qquad\qquad L_\mathrm{r} = 9.06 \text{ h}, \\
G_i(0) &= \begin{cases} 18.10 & \text{for } i = 1, \ldots, 5, \\ 7.51 & \text{for } i = 6, \ldots, 10, \end{cases} \qquad
\begin{aligned} L_\mathrm{y} &= 6.50 \text{ h}, \\ L_\mathrm{g} &= 15.79 \text{ h}. \end{aligned}
\end{aligned}
\tag{S18}
$$

Note that $L_\mathrm{r} + L_\mathrm{y} + L_\mathrm{g} = 31.35$ h, in good agreement with the observed cell cycle time of 27 h.

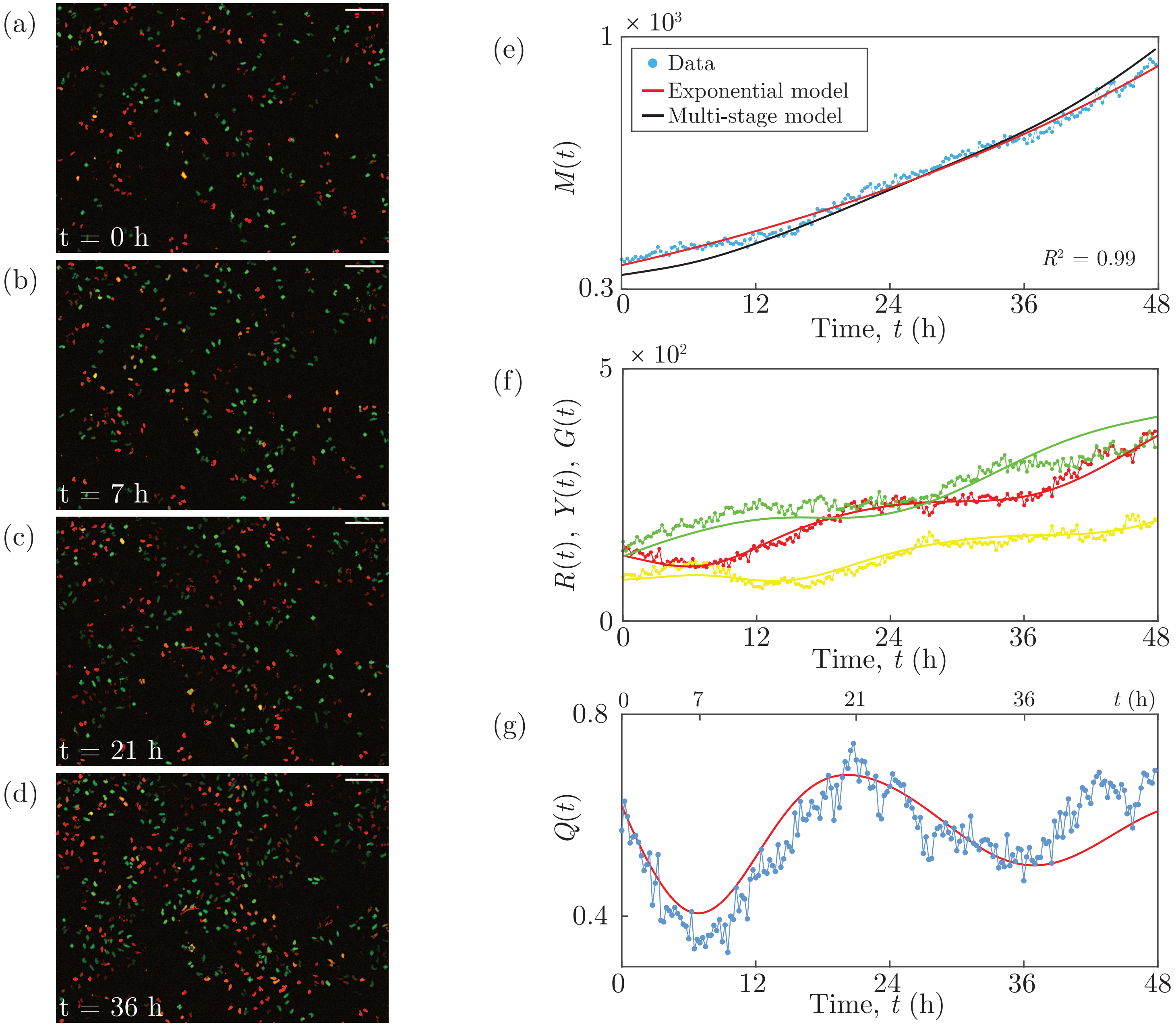

**Figure S5: WM983C experimental data and multi-stage model solution.** (a)–(d) Images of a proliferation assay with FUCCI-WM983C cells. Scale bar 200 $\mu$m. (e) $M(t)$. Linear regression of $\ln M(t)$ versus $t$ provides $R^2$. (f) $R(t)$, $Y(t)$ and $G(t)$. (g) $Q(t)$. Experimental data are shown as discs and the model solutions as curves.

###### 4.2.7 WM983C cell line - Figure S6

The experimentally-determined mean cell cycle time for WM983C is $\mathcal{T} = 27$ h [10]. We partition each cell cycle phase into $N = 20$ stages, giving a total of $k = 60$ stages for the complete cell cycle. From the start of each phase we set every 10 successive stages to have equal numbers of cells. We therefore only require a total of 6 distinct population parameters.

The vector objective function is $\mathbf{F}(\mathbf{x}) = [\,\mathbf{f}_1(\mathbf{x})\;\; 10^{-3}\mathbf{f}_2(\mathbf{x})\;\; 10^{-3}\mathbf{f}_3(\mathbf{x})\;\; 10^{-3}\mathbf{f}_4(\mathbf{x})\;\; 10^{-2}\mathbf{f}_5(\mathbf{x})\,]$. Trialling different parameters chosen randomly and uniformly from $(0.1, 1)$ for $R_i(0)$, $Y_i(0)$ and $G_i(0)$, and from $(4, 20)$ for $L_\mathrm{r}$, $L_\mathrm{y}$ and $L_\mathrm{g}$, we obtain the parameterisation

$$R_i(0) = \begin{cases} 3.47 & \text{for } i = 1, \ldots, 10, \\ 2.35 & \text{for } i = 11, \ldots, 20, \end{cases} \qquad Y_i(0) = \begin{cases} 1.10 & \text{for } i = 1, \ldots, 10, \\ 2.09 & \text{for } i = 10, \ldots, 20, \end{cases}$$

$$L_\mathrm{r} = 9.22 \text{ h}, \tag{S19}$$

$$G_i(0) = \begin{cases} 2.68 & \text{for } i = 1, \ldots, 10, \\ 1.64 & \text{for } i = 11, \ldots, 20, \end{cases} \qquad \begin{aligned} L_\mathrm{y} &= 5.46 \text{ h}, \\ L_\mathrm{g} &= 14.44 \text{ h}. \end{aligned}$$

Note that $L_\mathrm{r} + L_\mathrm{y} + L_\mathrm{g} = 29.12$ h, in good agreement with the observed cell cycle time of 27 h.

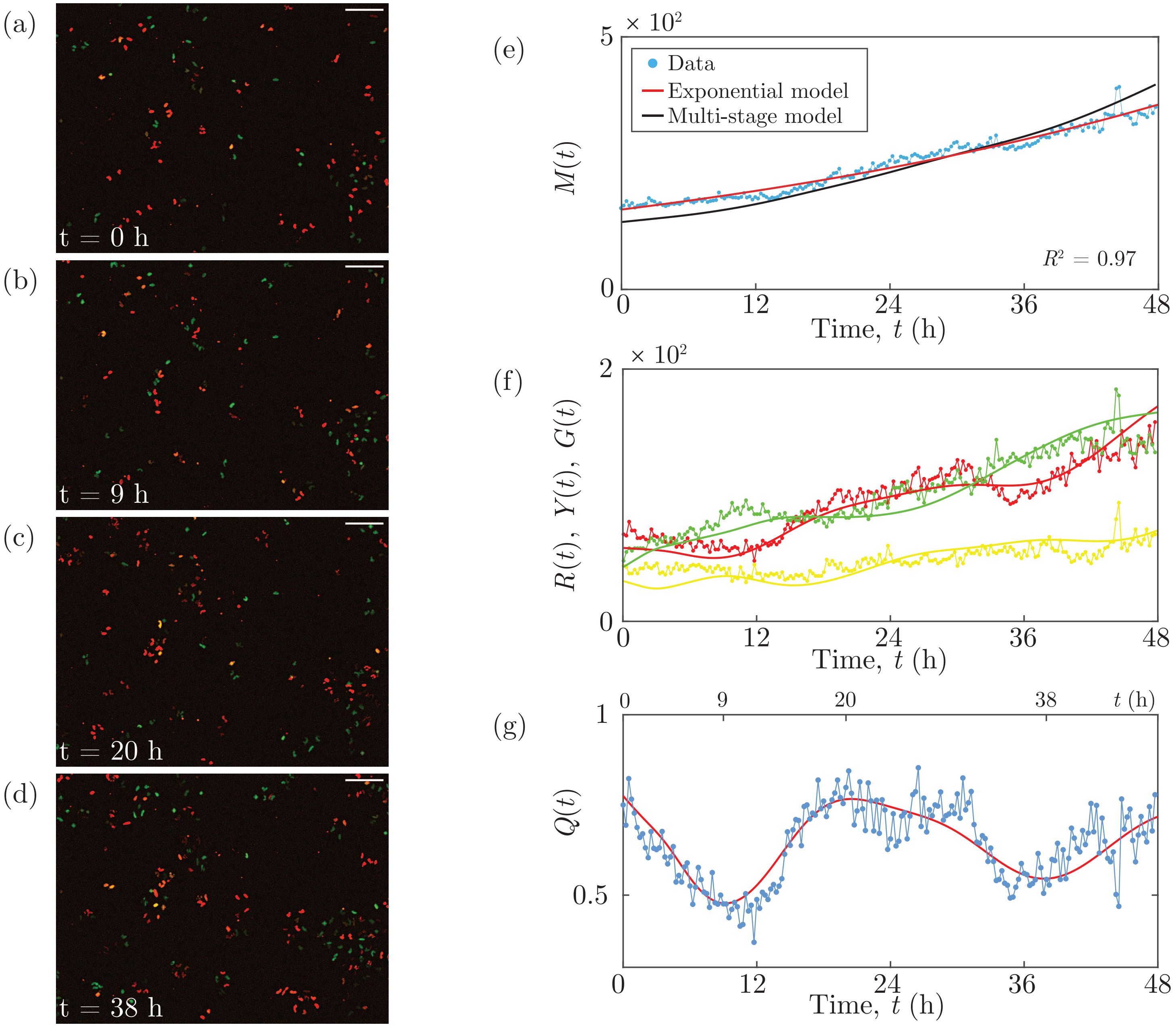

**Figure S6: WM983C experimental data and multi-stage model solution.** (a)–(d) Images of a proliferation assay with FUCCI-WM983C cells. Scale bar 200 $\mu$m. (e) $M(t)$. Linear regression of $\ln M(t)$ versus $t$ provides $R^2$. (f) $R(t)$, $Y(t)$ and $G(t)$. (g) $Q(t)$. Experimental data are shown as discs and the model solutions as curves.

###### 4.2.8 1205Lu cell line - Figure S7

The experimentally-determined mean cell cycle time for 1205Lu is $\mathcal{T} = 36$ h [10]. We partition each cell cycle phase into $N = 20$ stages, giving a total of $k = 60$ stages for the complete cell cycle. From the start of each phase we set every 5 successive stages to have equal numbers of cells. We therefore only require a total of 12 distinct population parameters.

The vector objective function is $\mathbf{F}(\mathbf{x}) = [\,\mathbf{f}_1(\mathbf{x})\;\; 10^{-2}\mathbf{f}_2(\mathbf{x})\;\; 10^{-2}\mathbf{f}_3(\mathbf{x})\;\; 10^{-2}\mathbf{f}_4(\mathbf{x})\;\; 0.5\,\mathbf{f}_6(\mathbf{x})\,]$. Starting with the parameters

$$
\begin{aligned}
R_i(0) &= \begin{cases} 0.27 \quad \text{for } i = 1, \dots, 5, \\ 0 \quad \text{for } i = 6, \dots, 10, \\ 21.78 \quad \text{for } i = 11, \dots, 15, \\ 0 \quad \text{for } i = 16, \dots, 20, \end{cases} \qquad
Y_i(0) = \begin{cases} 4.73 \quad \text{for } i = 1, \dots, 5, \\ 5.39 \quad \text{for } i = 6, \dots, 10, \\ 0 \quad \text{for } i = 11, \dots, 15, \\ 2.49 \quad \text{for } i = 16, \dots, 20, \end{cases} \\
G_i(0) &= \begin{cases} 3.15 \quad \text{for } i = 1, \dots, 5, \\ 5.17 \quad \text{for } i = 6, \dots, 10, \\ 2.08 \quad \text{for } i = 11, \dots, 15, \\ 0.45 \quad \text{for } i = 16, \dots, 20, \end{cases} \qquad
\begin{aligned} L_{\mathrm{r}} &= 20.97 \text{ h}, \\ L_{\mathrm{y}} &= 10.07 \text{ h}, \\ L_{\mathrm{g}} &= 10.52 \text{ h}, \end{aligned}
\end{aligned} \tag{S20}
$$

we obtain the parameterisation

$$
R_i(0) = \begin{cases} 0 \quad \text{for } i = 1, \ldots, 5, \\ 13.79 \quad \text{for } i = 6, \ldots, 10, \\ 5.73 \quad \text{for } i = 11, \ldots, 15, \\ 4.19 \quad \text{for } i = 16, \ldots, 20, \end{cases} \qquad Y_i(0) = \begin{cases} 9.39 \quad \text{for } i = 1, \ldots, 5, \\ 0 \quad \text{for } i = 6, \ldots, 10, \\ 0 \quad \text{for } i = 11, \ldots, 15, \\ 5.04 \quad \text{for } i = 16, \ldots, 20, \end{cases}
$$

$$
G_i(0) = \begin{cases} 2.31 \quad \text{for } i = 1, \ldots, 5, \\ 0 \quad \text{for } i = 6, \ldots, 10, \\ 3.52 \quad \text{for } i = 11, \ldots, 15, \\ 0.63 \quad \text{for } i = 16, \ldots, 20, \end{cases} \qquad \begin{aligned} L_{\mathrm{r}} &= 19.61 \text{ h}, \\ L_{\mathrm{y}} &= 7.99 \text{ h}, \\ L_{\mathrm{g}} &= 10.76 \text{ h}, \end{aligned} \tag{S21}
$$

Note that $L_{\mathrm{r}} + L_{\mathrm{y}} + L_{\mathrm{g}} = 38.36$ h, in good agreement with the observed cell cycle time of 36 h.

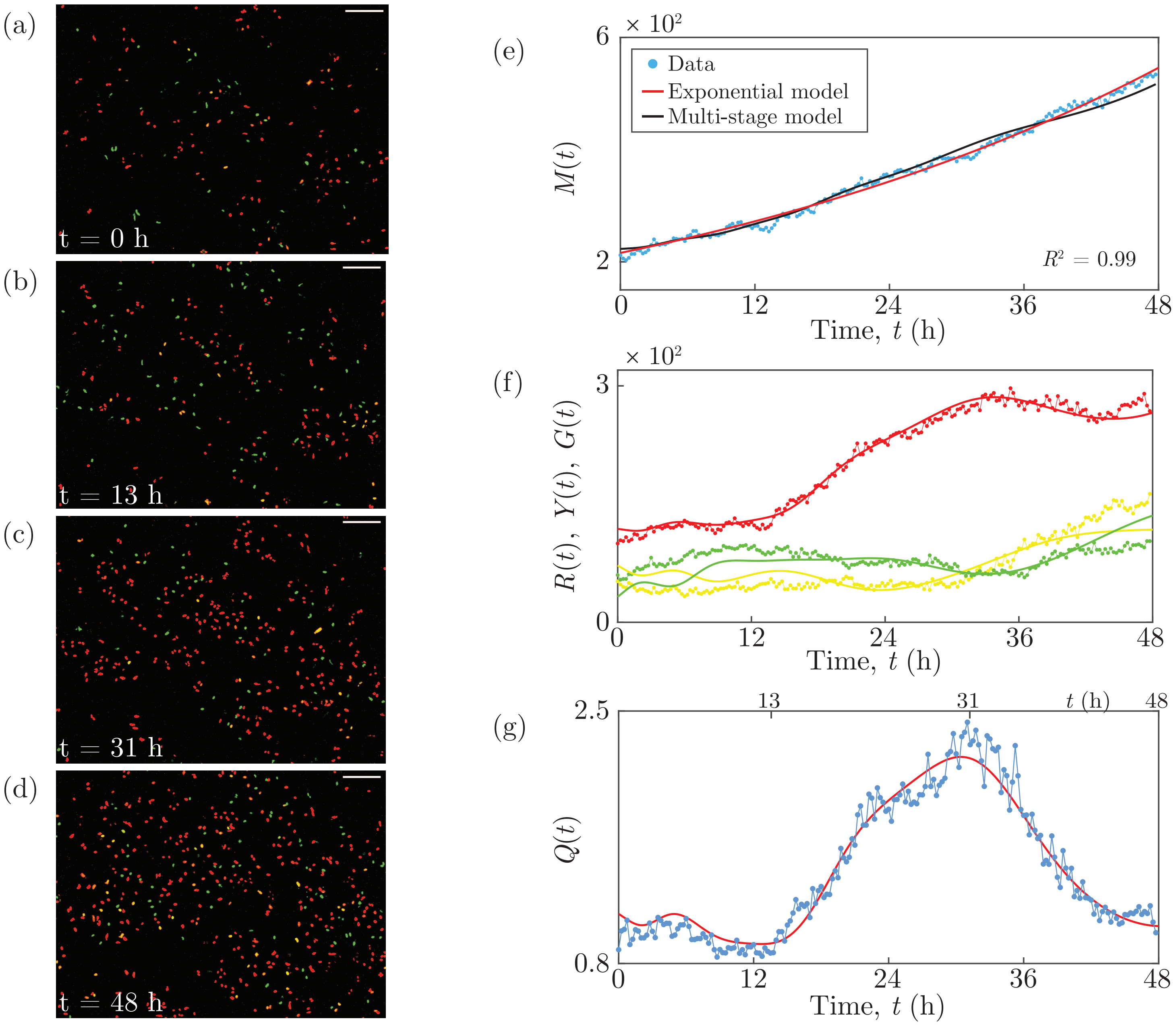

**Figure S7: 1205Lu experimental data and multi-stage model solution.** (a)–(d) Images of a proliferation assay with FUCCI-1205Lu cells. Scale bar 200 $\mu$m. (e) $M(t)$. Linear regression of $\ln M(t)$ versus $t$ provides $R^2$. (f) $R(t)$, $Y(t)$ and $G(t)$. (g) $Q(t)$. Experimental data are shown as discs and the model solutions as curves.

###### 4.2.9 1205Lu cell line - Figure S8

The experimentally-determined mean cell cycle time for 1205Lu is $\mathcal{T} = 36$ h [10]. We partition each cell cycle phase into $N = 20$ stages, giving a total of $k = 60$ stages for the complete cell cycle. From the start of each phase we set every 4 successive stages to have equal numbers of cells. We therefore only require a total of 15 distinct population parameters.

The vector objective function is $\mathbf{F}(\mathbf{x}) = [\,\mathbf{f}_1(\mathbf{x})\;\; 10^{-2}\mathbf{f}_2(\mathbf{x})\;\; 10^{-2}\mathbf{f}_3(\mathbf{x})\;\; 10^{-2}\mathbf{f}_4(\mathbf{x})\;\; 0.6\,\mathbf{f}_6(\mathbf{x})\,]$. Trialling different parameters chosen randomly and uniformly from $(0.1, 1)$ for $R_i(0)$, $Y_i(0)$ and $G_i(0)$, and from $(4, 20)$ for $L_\mathrm{r}$, $L_\mathrm{y}$ and $L_\mathrm{g}$, we obtain the parameterisation

$$R_i(0) = \begin{cases} 0 \quad \text{for } i = 1, \ldots, 4, \\ 23.75 \quad \text{for } i = 5, \ldots, 8, \\ 0 \quad \text{for } i = 9, \ldots, 12, \\ 13.54 \quad \text{for } i = 13, \ldots, 16, \\ 0.92 \quad \text{for } i = 17, \ldots, 20, \end{cases} \qquad Y_i(0) = \begin{cases} 12.55 \quad \text{for } i = 1, \ldots, 4, \\ 0 \quad \text{for } i = 5, \ldots, 8, \\ 0 \quad \text{for } i = 9, \ldots, 12, \\ 0 \quad \text{for } i = 13, \ldots, 16, \\ 0.67 \quad \text{for } i = 17, \ldots, 20, \end{cases} \tag{S21}$$

$$G_i(0) = \begin{cases} 9.94 \quad \text{for } i = 1, \ldots, 4, \\ 0 \quad \text{for } i = 5, \ldots, 8, \\ 3.24 \quad \text{for } i = 9, \ldots, 12, \\ 1.46 \quad \text{for } i = 13, \ldots, 16, \\ 0 \quad \text{for } i = 17, \ldots, 20, \end{cases} \qquad \begin{aligned} L_\mathrm{r} &= 19.20 \text{ h}, \\ L_\mathrm{y} &= 7.98 \text{ h}, \\ L_\mathrm{g} &= 10.89 \text{ h}, \end{aligned}$$

Note that $L_\mathrm{r} + L_\mathrm{y} + L_\mathrm{g} = 38.07$ h, in good agreement with the observed cell cycle time of 36 h.

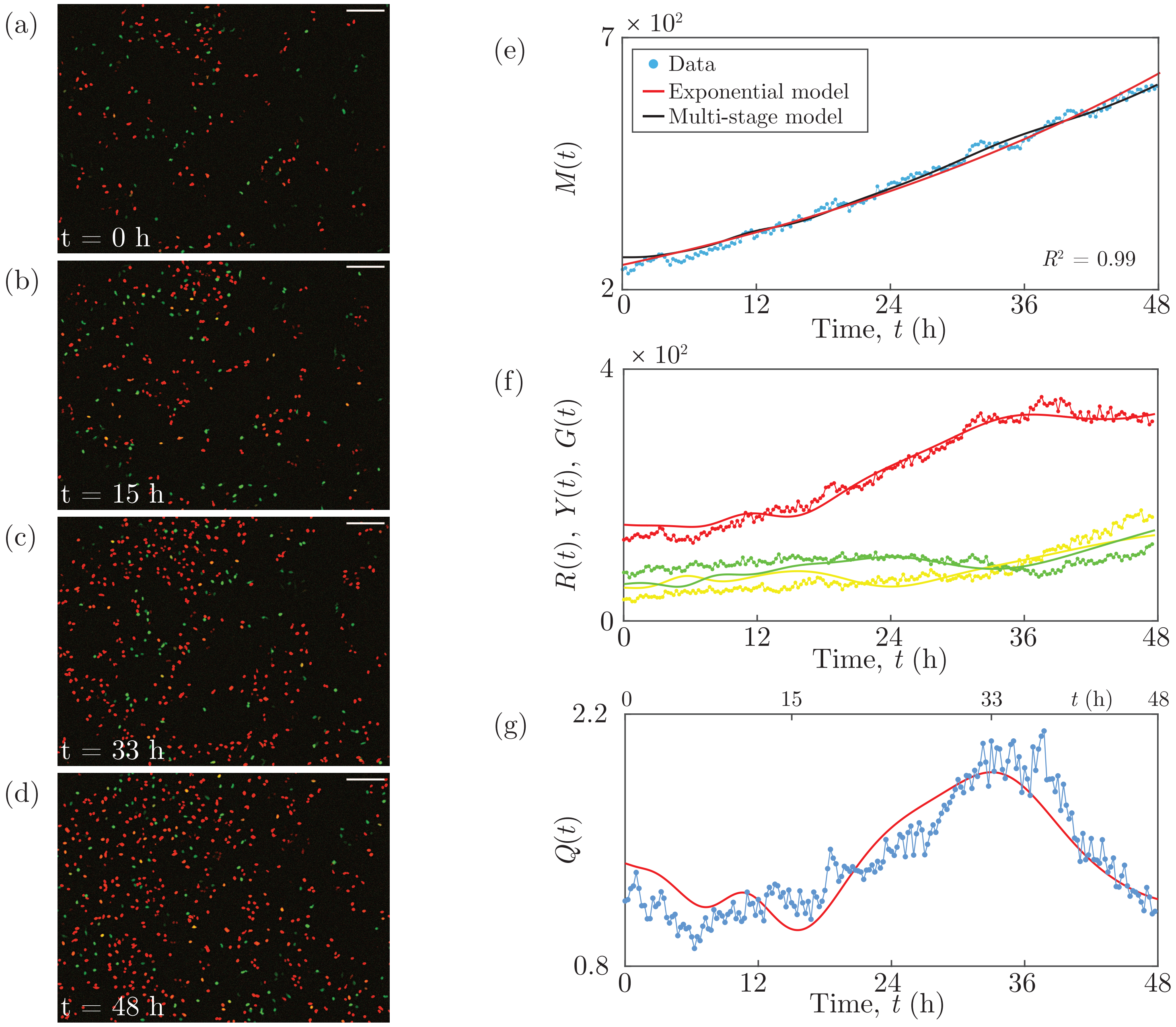

**Figure S8: 1205Lu experimental data and multi-stage model solution.** (a)–(d) Images of a proliferation assay with FUCCI-1205Lu cells. Scale bar 200 $\mu$m. (e) $M(t)$. Linear regression of $\ln M(t)$ versus $t$ provides $R^2$. (f) $R(t)$, $Y(t)$ and $G(t)$. (g) $Q(t)$. Experimental data are shown as discs and the model solutions as curves.

###### 4.2.10 1205Lu cell line - Figure S9

The experimentally-determined mean cell cycle time for 1205Lu is $\mathcal{T} = 36$ h [10]. We partition each cell cycle phase into $N = 12$ stages, giving a total of $k = 36$ stages for the complete cell cycle. From the start of each phase we set every 3 successive stages to have equal numbers of cells. We therefore only require a total of 12 distinct population parameters.

The vector objective function is $\mathbf{F}(\mathbf{x}) = [\,\mathbf{f}_1(\mathbf{x})\;\; 3\times10^{-3}\mathbf{f}_2(\mathbf{x})\;\; 3\times10^{-3}\mathbf{f}_3(\mathbf{x})\;\; 3\times10^{-3}\mathbf{f}_4(\mathbf{x})\;\; 0.2\,\mathbf{f}_6(\mathbf{x})\,]$. Trialling different parameters chosen randomly and uniformly from $(0.1, 1)$ for $R_i(0)$, $Y_i(0)$ and $G_i(0)$, and from $(4, 20)$ for $L_\mathrm{r}$, $L_\mathrm{y}$ and $L_\mathrm{g}$, we obtain the parameterisation

$$
\begin{aligned}
R_i(0) &= \begin{cases} 0 \quad \text{for } i = 1,\dots,3, \\ 24.89 \quad \text{for } i = 4,\dots,6, \\ 25.23 \quad \text{for } i = 7,\dots,9, \\ 0 \quad \text{for } i = 10,\dots,12, \end{cases}
\qquad
Y_i(0) = \begin{cases} 20.14 \quad \text{for } i = 1,\dots,3, \\ 0 \quad \text{for } i = 4,\dots,6, \\ 0 \quad \text{for } i = 7,\dots,9, \\ 0 \quad \text{for } i = 10,\dots,12, \end{cases} \\
G_i(0) &= \begin{cases} 2.68 \quad \text{for } i = 1,\dots,3, \\ 20.35 \quad \text{for } i = 4,\dots,6, \\ 0 \quad \text{for } i = 7,\dots,9, \\ 0 \quad \text{for } i = 10,\dots,12, \end{cases}
\qquad
\begin{aligned} L_\mathrm{r} &= 19.18 \text{ h}, \\ L_\mathrm{y} &= 9.08 \text{ h}, \\ L_\mathrm{g} &= 10.99 \text{ h}, \end{aligned}
\end{aligned}
\tag{S22}
$$

Note that $L_\mathrm{r} + L_\mathrm{y} + L_\mathrm{g} = 39.25$ h, in good agreement with the observed cell cycle time of 36 h.

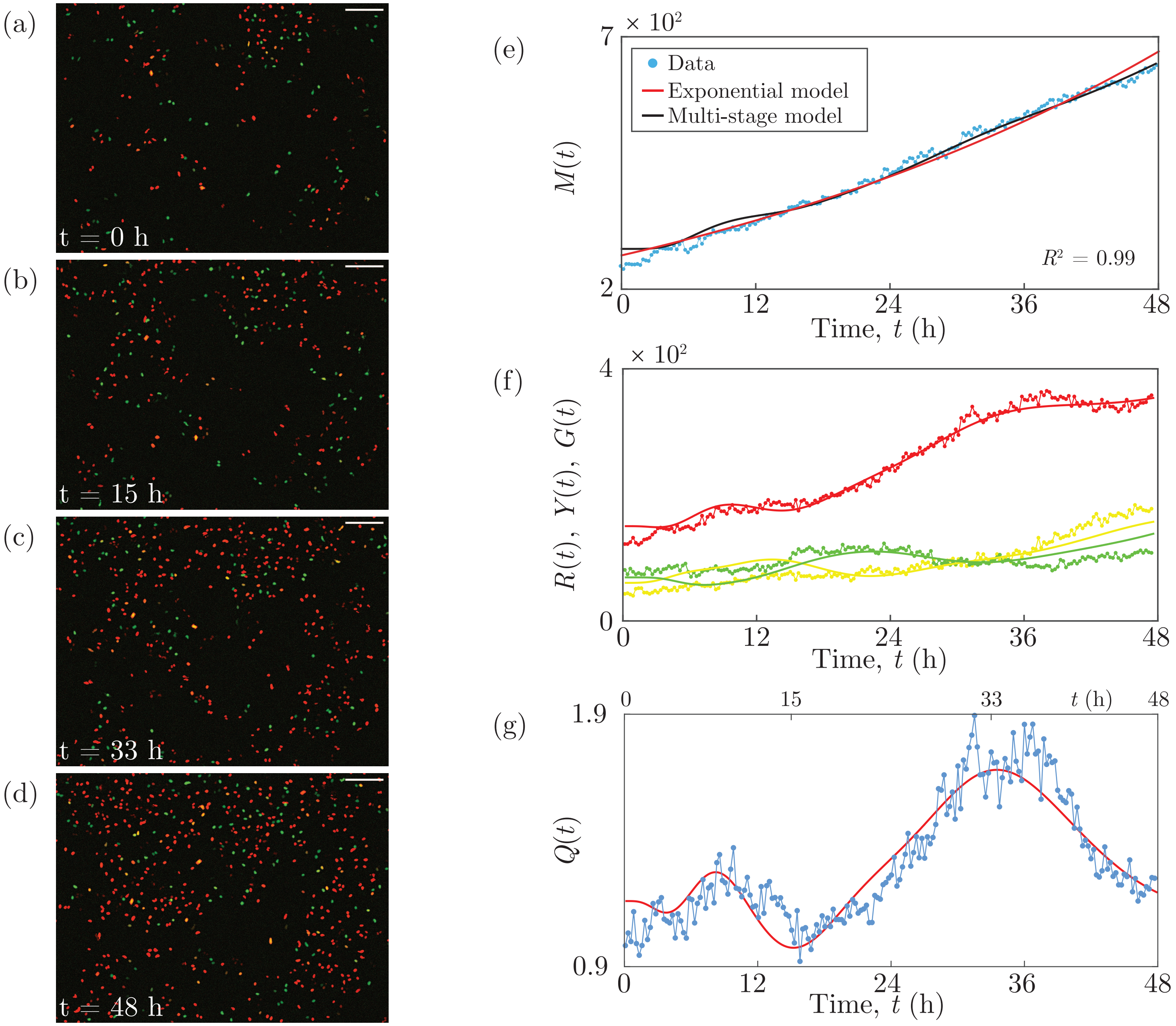

**Figure S9: 1205Lu experimental data and multi-stage model solution.** (a)–(d) Images of a proliferation assay with FUCCI-1205Lu cells. Scale bar 200 $\mu$m. (e) $M(t)$. Linear regression of $\ln M(t)$ versus $t$ provides $R^2$. (f) $R(t)$, $Y(t)$ and $G(t)$. (g) $Q(t)$. Experimental data are shown as discs and the model solutions as curves.

### 5 All experimental data

Here we provide all of our new experimental data in the following forms:

- The total number of cells $M(t)$;
- The ratio $Q(t)$ of the number of cells in G1 to the combined number of cells in eS and S/G2/M;
- The discrete Fourier transform of $Q(t)$.

These data are obtained from the three cell lines C8161, WM983C and 1205Lu, and four independent experiments. In Experiments 1–3 we use one well of a 24-well plate, and in Experiment 4 we use two wells of a 24-well plate. From each well we obtain time-series stacks at six different positions. These data are available in Supporting Information 2–4, in the form of the number of cells in each phase, G1, eS and S/G2/M, at each time point.

In every experiment, the population growth $M(t)$ appears to be exponential, and the ratio $Q(t)$ reveals the presence of inherent synchronisation. In a given well, the six different positions can exhibit different degrees of inherent synchronisation. Further, the synchronisation can be out of phase between the different positions in a given well, and between the different wells.

Note that for some of the data there are a couple of consecutive time points which show a much higher total number of cells than expected, and a corresponding lower ratio in the ratio data. This is due to a large decrease in the signal-to-noise ratio in the green channel at these time points. The specific cause of this is unknown, however fluorescence microscopy is subject to such variations in the signal-to-noise ratio at times. As there is such a large reduction in the signal-to-noise ratio, it is not possible to reduce the unwanted noise without compromising the signal quality.

We provide the discrete Fourier transforms of $Q(t)$ for every data set to quantitatively demonstrate the existence of oscillating subpopulations in our experimental data. The transforms are obtained using the fast Fourier transform `fft` function [16] in MATLAB, without spectral interpolation. For clarity, we would like the amplitude of the Fourier transform to be zero at zero frequency, so we apply the transform to the time series $Q(t) - \overline{Q(t)}$, where $\overline{Q(t)}$ is the mean value of the time series. The transformed data are presented as single-sided spectra showing the magnitude of the Fourier transform, $A(f)$, as a function of frequency, $f$, where $0 \text{ h}^{-1} \leq f \leq 2 \text{ h}^{-1}$. Note that the Nyquist frequency is $2 \text{ h}^{-1}$.

The Fourier transforms all show dominant frequencies corresponding to periods of either 16, 24 or 48 h, which clearly indicate the presence of oscillations in $Q(t)$ for each of our data sets, in accordance with the existence of inherent synchronisation. These periods are a reasonable approximation of the experimental cell-cycle durations of the cell lines. To increase the resolution of the frequencies, and thereby obtain better estimates of the periods of the oscillations, a time interval between the time-series images which is less than 15 minutes is required.

#### 5.1 C8161 cell line

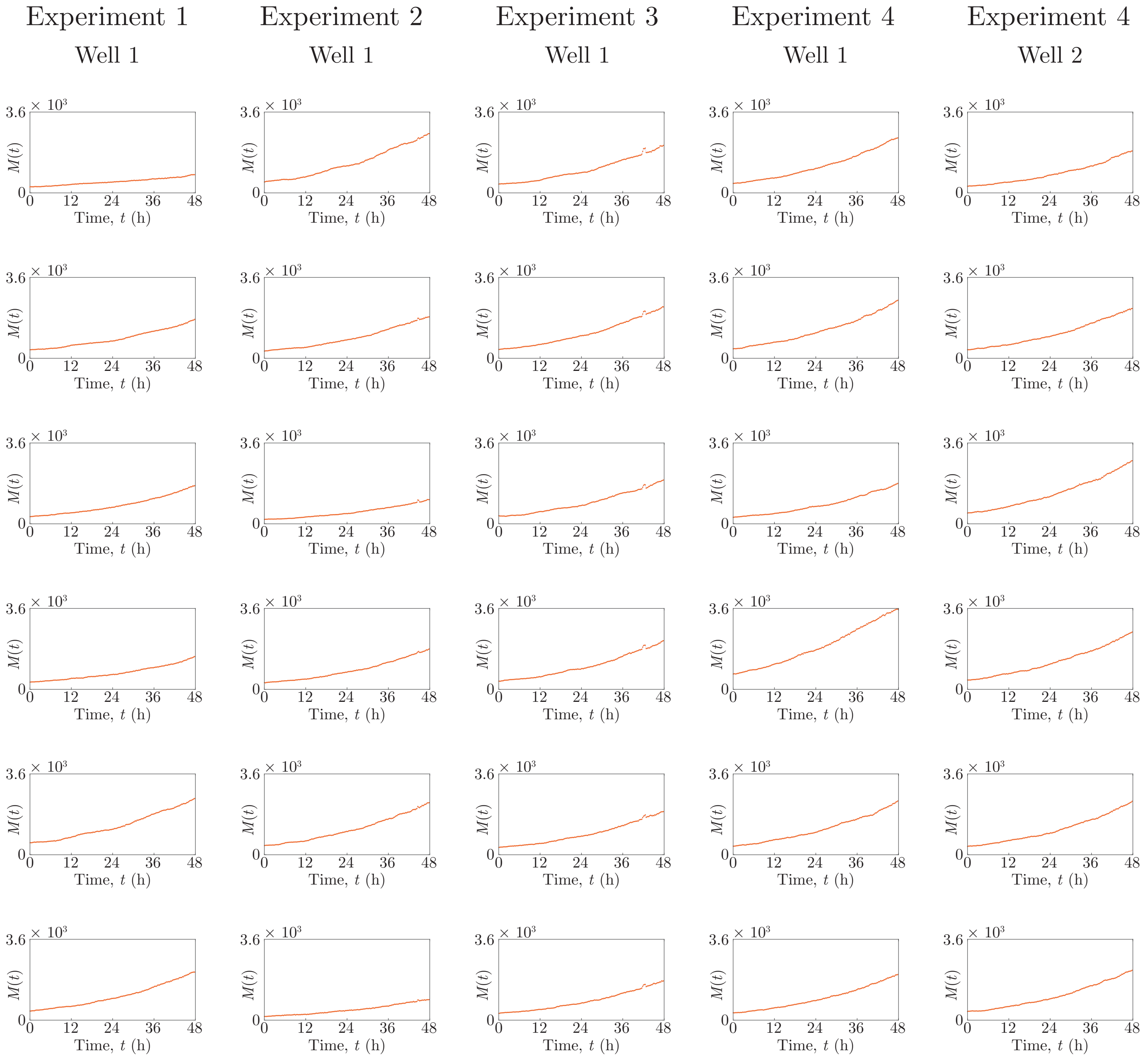

**Figure S10: C8161 experimental data.** Total number of cells $M(t)$.

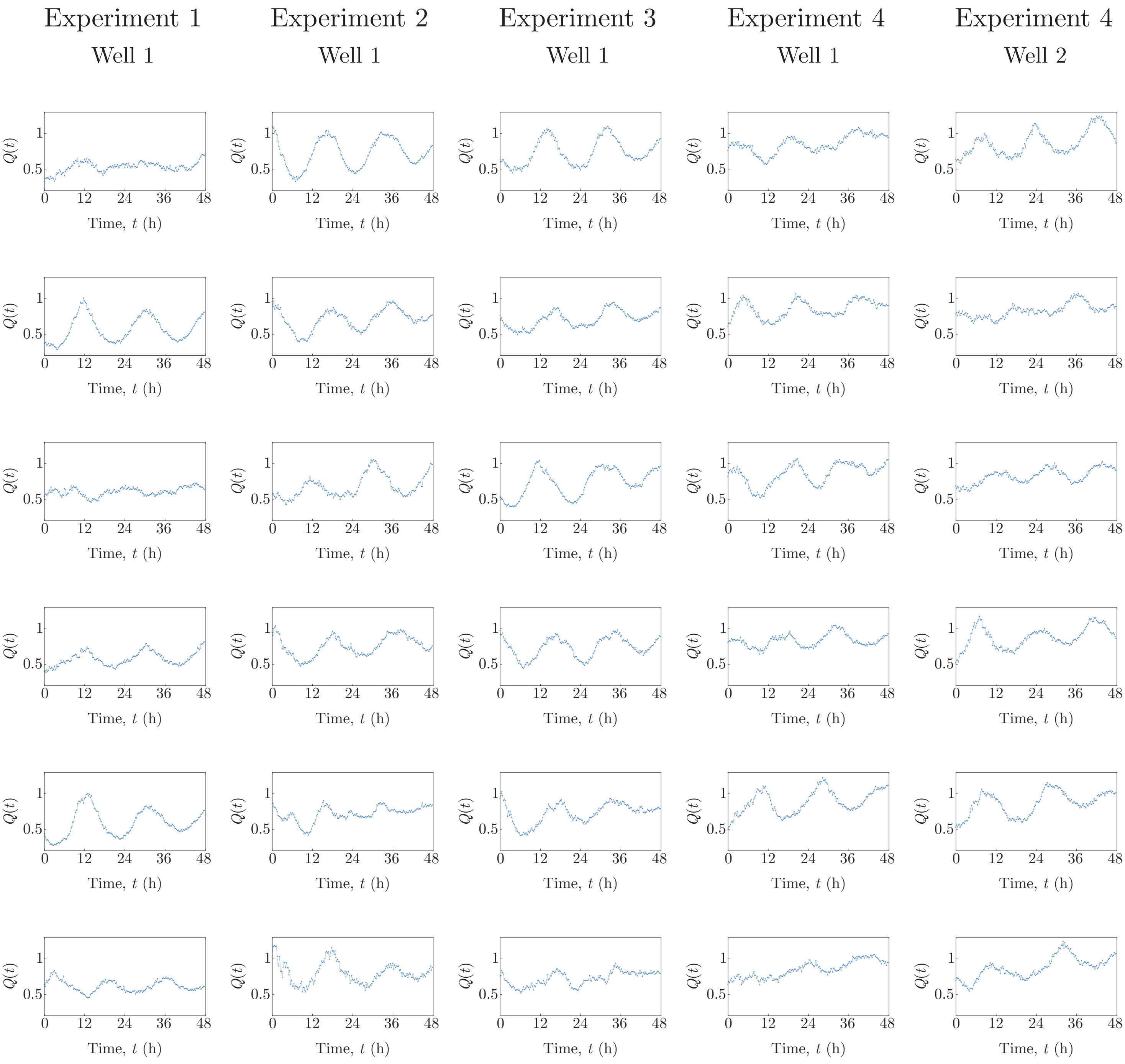

**Figure S11: C8161 experimental data.** Ratio $Q(t)$ of the number of cells in G1 to the number of cells in eS and S/G2/M.

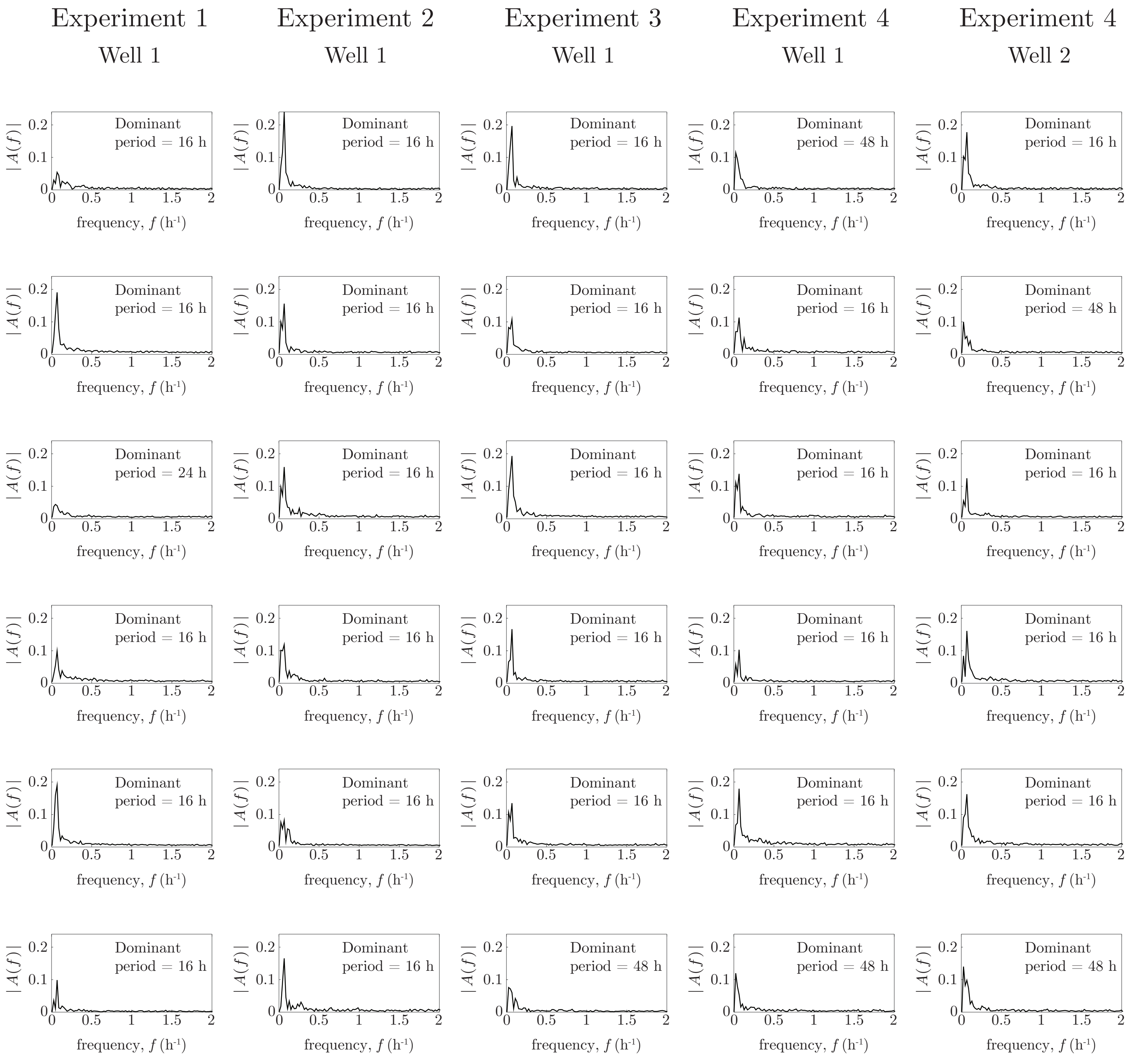

**Figure S12: C8161 experimental data.** Magnitude of the Fourier transform, $A(f)$, of the ratio $Q(t) - \overline{Q(t)}$, as a function of frequency, $f$.

#### 5.2 WM983C cell line

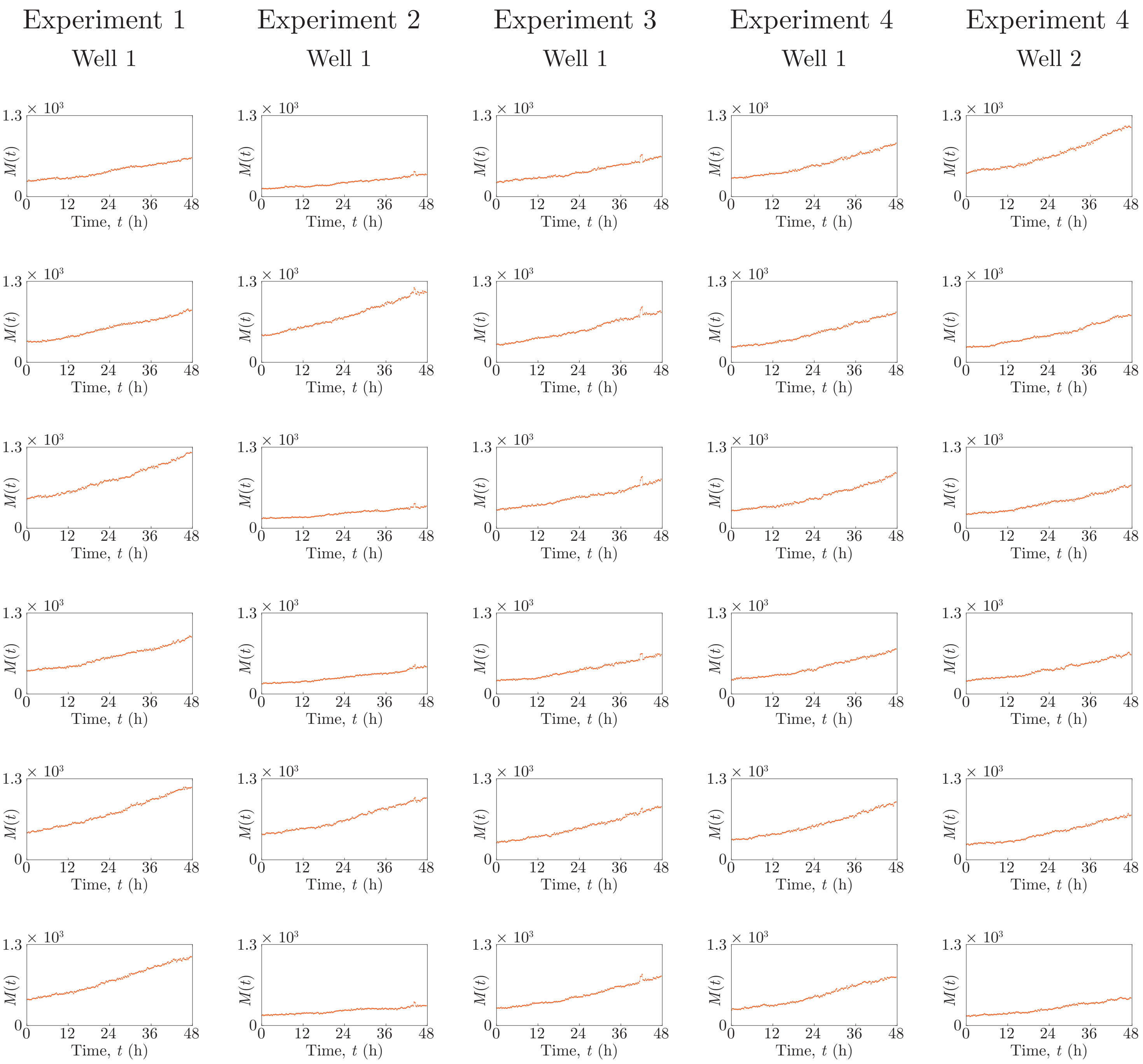

**Figure S13: WM983C experimental data.** Total number of cells $M(t)$.

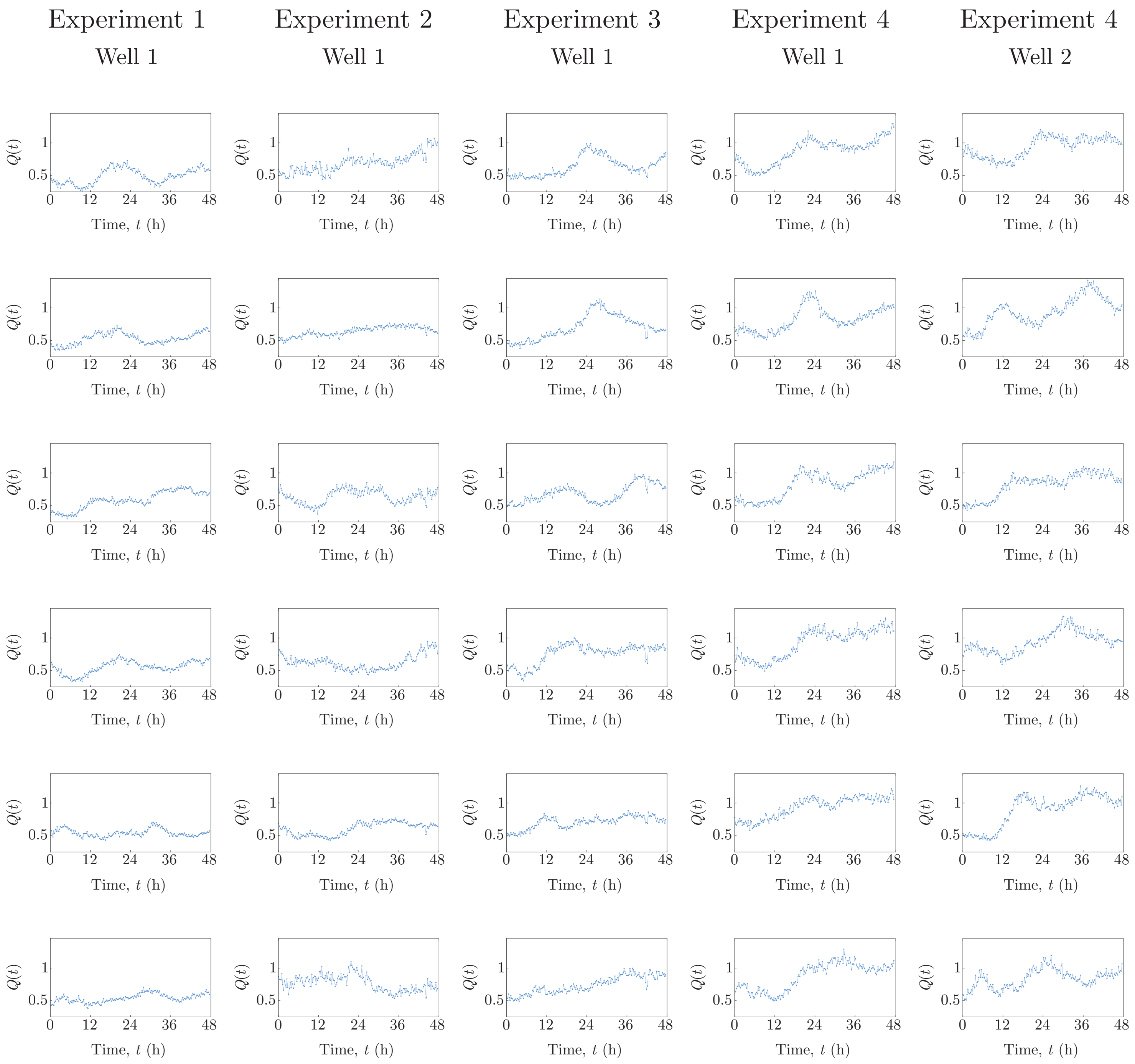

**Figure S14: WM983C experimental data.** Ratio $Q(t)$ of the number of cells in G1 to the number of cells in eS and S/G2/M.

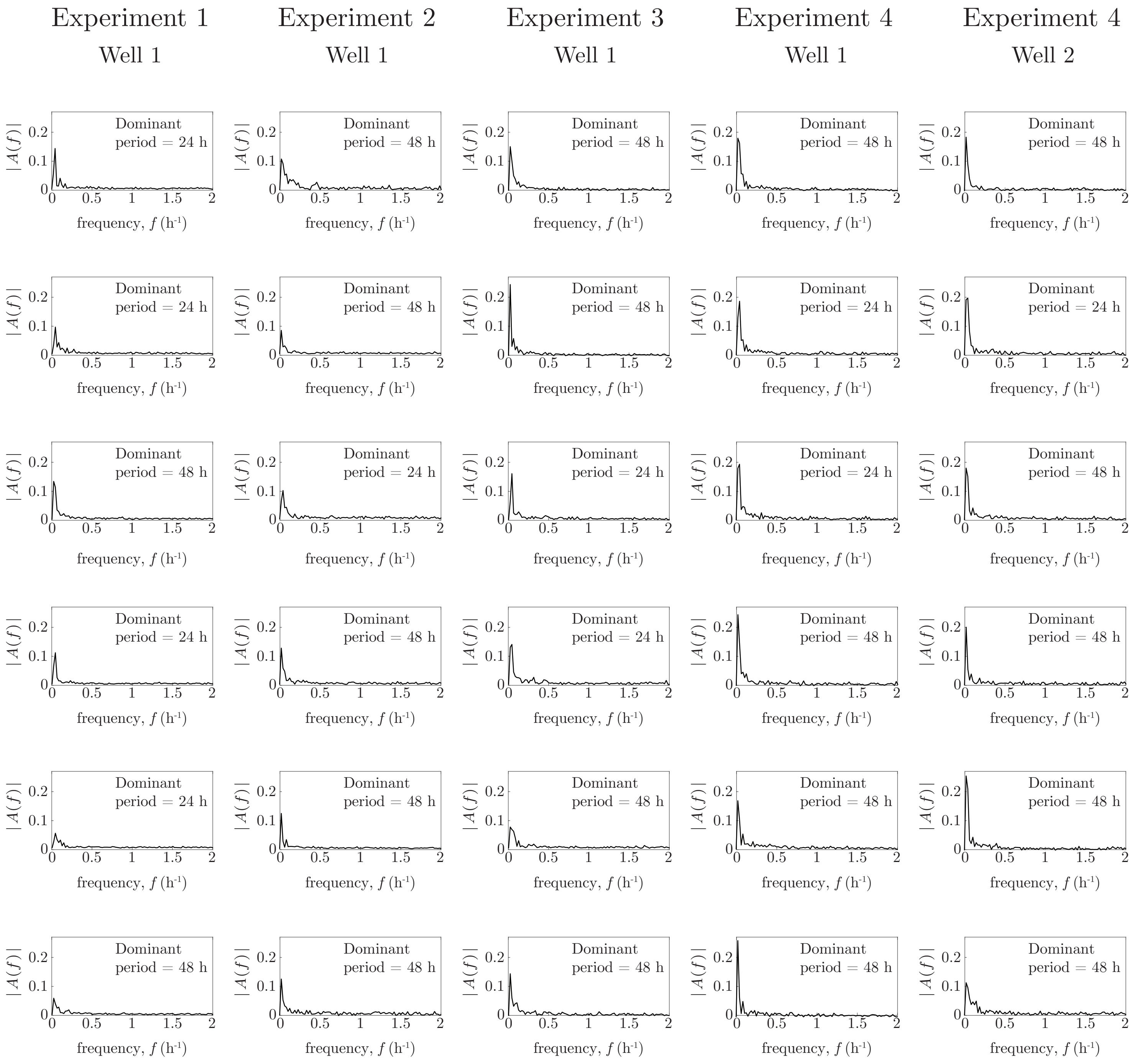

**Figure S15: WM983C experimental data.** Magnitude of the Fourier transform, $A(f)$, of the ratio $Q(t) - \overline{Q(t)}$, as a function of frequency, $f$.

#### 5.3 1205Lu cell line

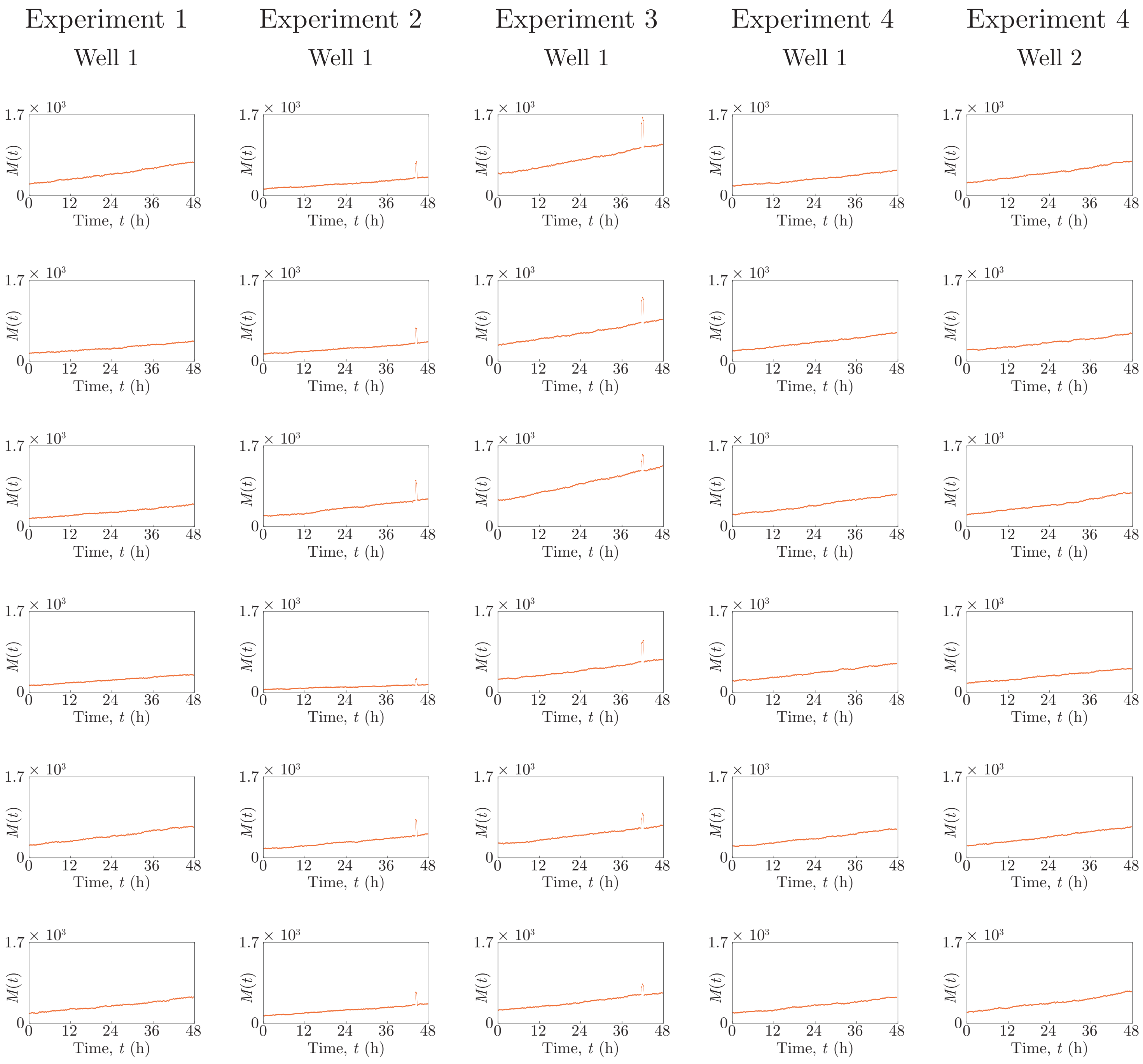

**Figure S16: 1205Lu experimental data.** Total number of cells $M(t)$.

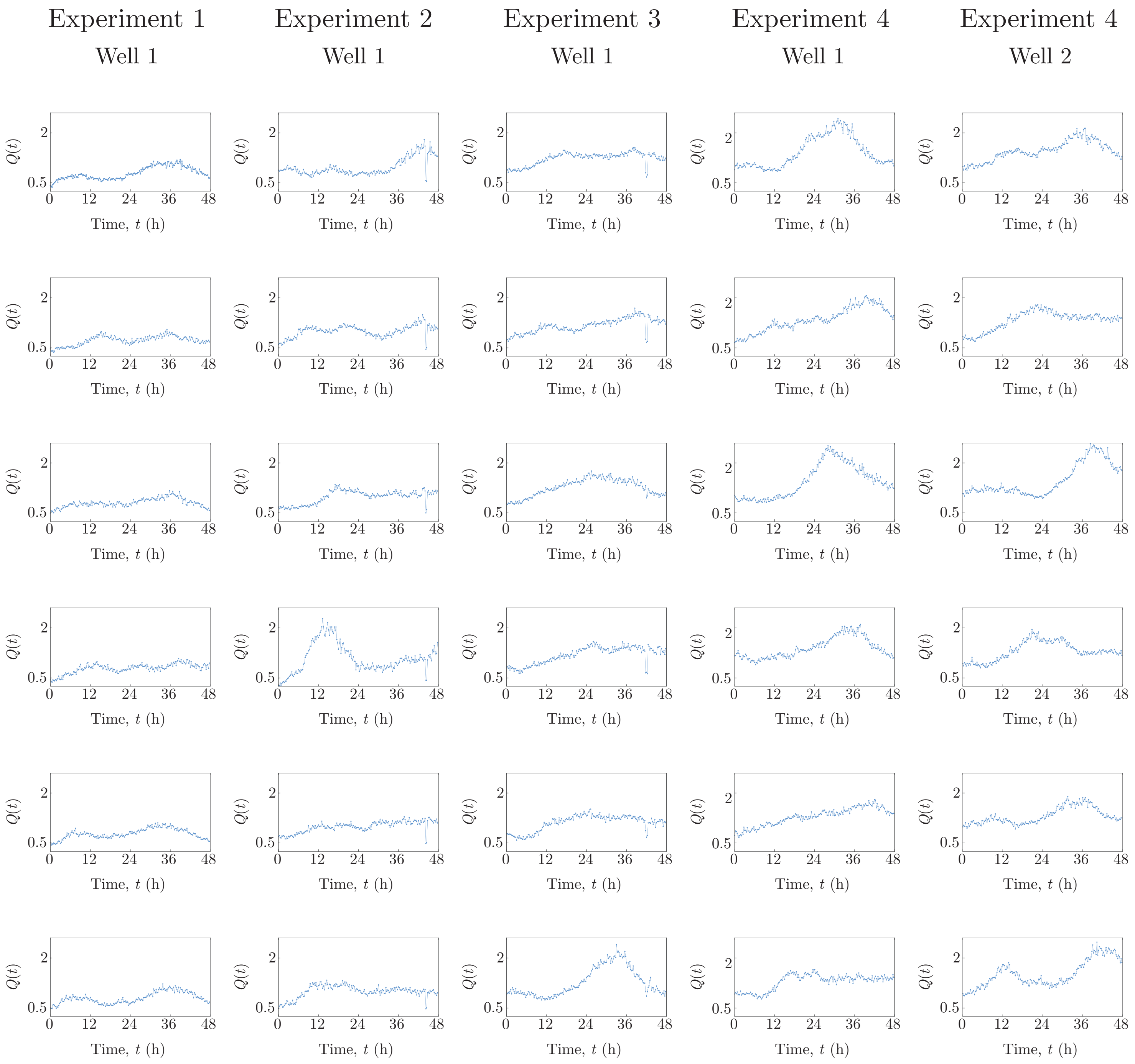

**Figure S17: 1205Lu experimental data.** Ratio $Q(t)$ of the number of cells in G1 to the number of cells in eS and S/G2/M.

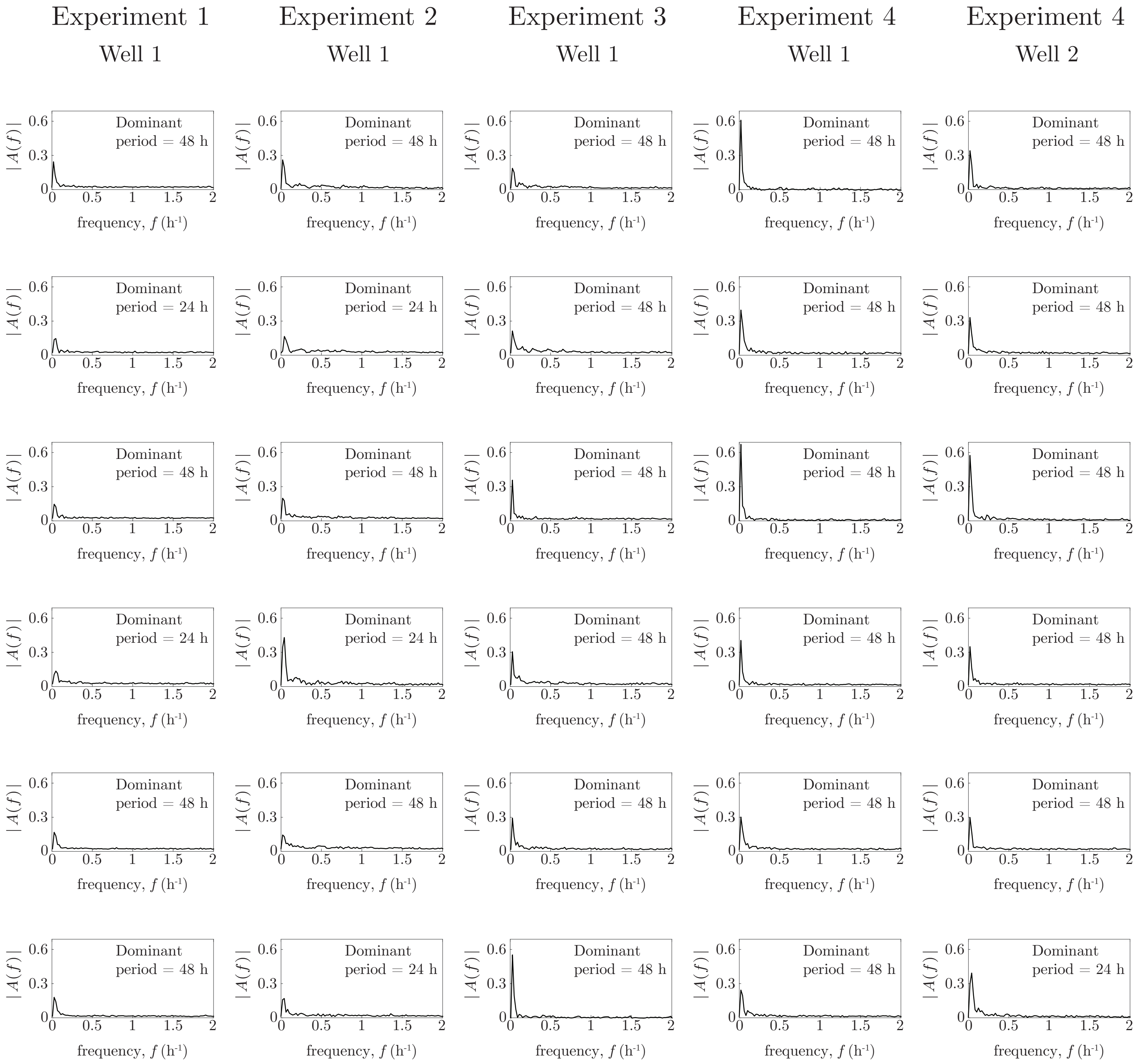

**Figure S18: 1205Lu experimental data.** Magnitude of the Fourier transform, $A(f)$, of the ratio $Q(t) - \overline{Q(t)}$, as a function of frequency, $f$.
